# Loss of canonical sex steroid signaling correlates with prostate cancer progression and induces tumor escape in *Drosophila*

**DOI:** 10.1101/2025.03.18.643888

**Authors:** M. Vialat, E. Baabdaty, C. Vachias, F. Degoul, A. Trousson, P. Pouchin, JMA. Lobaccaro, S. Baron, L. Morel, C. de Joussineau

**Affiliations:** Université Clermont Auvergne, CNRS 6293, INSERM 1103, iGReD; Clermont–Ferrand, France; Groupe Cancer Clermont Auvergne; Clermont-Ferrand, France

## Abstract

In cancer, tumor escape often arises following treatments. This is especially true during deprivation therapies, where this phenomenon has been linked to steroid signaling reactivation despite deprivation. Here, we show that in prostate cancer tissues the canonical androgen pathway itself is deactivated and, in fact, loss of canonical AR signaling tightly correlates with cancer progression. This raises the possibility that loss of canonical sex steroid signaling could promote the progression. We tested this hypothesis a drosophila model of prostate cancer. There, repression of canonical sex steroid ecdysone receptor signaling displays both anti- and protumor effects. On the one hand, it slightly decreases extra-epithelial tumor formation, and increases their propensity for apoptosis. On the other hand, it induces the growth of a new tumor cell population, which growth is normally prevented by autocrine/intracrine ecdysone signaling. Furthermore, this population appears to emerge from epithelial clones within the gland. To do so, tumor cells change the way they migrate out of the epithelium, forming a new layer between the normal epithelium and the basement membrane. This depends on a modification of epithelial basal extrusion that likely relies on the downregulation of Ecdysone target gene *αTub60D*. Thus, in drosophila accessory gland, lack of sex steroid signaling not only coincides with but actually induces tumor escape. Together, these results question the role of sex steroid deprivation on tumor progression, and point to altered basal extrusion as a possible mechanism shaping tumor escape.

## Introduction

Epithelial tumorigenesis is a complex, multistep process spanning from hypothetical initiation to metastatic spread. Late stages are better characterized, and have been associated with numerous alterations in molecular mechanisms that appear necessary for tumor progression (reviewed in (Hanahan and Weinberg, 2011)). Among them, specific pathways can act in specific tissues. Typically, the main sex steroids, estrogens and androgens, support growth of adenocarcinoma in reproductive tissues. Accordingly, endocrine therapy (sex steroid deprivation) is widely used in breast and prostate cancers to induce tumor regression.

However, despite strong initial response, tumor escape frequently occurs and even systematically in prostate cancer. Tumor escape is frequently associated with resistance to hormone deprivation. Treatment-resistant cells can adapt through high estrogen or androgen receptor (ER or AR) expression enabling sensitivity to reduced steroid levels, ER/AR mutations or alternative splicing leading to ligand-independent reactivation (Cao et al., 2014; Zundelevich et al., 2020). However, tumor escape also occurs when tumor cells become completely independent of steroid signaling for their progression, following *ER*/*AR* loss of function (Kuukasjärvi et al., 1996). So, it exists more than one path for tumor cells to survive endocrine therapy. Furthermore, cells that escape treatment appear more aggressive, with higher proliferative and metastatic potential (Clarke et al., 1989; Wu et al., 1994).

Despite best efforts, the origin of the cells that escape endocrine therapy remains debated, and the mechanisms allowing them to develop into tumors are still largely unknown. Over the last decades, cancer stem cells (CSCs) have emerged as strong candidates for initiating resistant tumor growth (Rodriguez et al., 2019; Rybak et al., 2015). Detected in many tumor types, these cells are defined by their insensitivity to treatments and their ability to recolonize a tumor that regresses upon treatment (Atashzar et al., 2020). Why these cells are present remains under scrutiny: hypothetically, they could already be present before treatment as a consequence of stochastic tumor heterogeneity, derive from adult stem cells, or transdifferentiate from tumor cells as a consequence of treatment selection. This last hypothesis, implying that deprivation therapy would in part be responsible for tumor progression, is supported by the observation that xenografted tumor cells can escape androgen deprivation in castrated mice (Iwasa et al., 2007; Nagabhushan et al., 1996; Wu et al., 1994; Zhou et al., 2004). However, the cellular and molecular mechanisms by which these cells could emerge and regrow to form a tumor are mostly unknown.

We previously developed a model of epithelial tumorigenesis in the drosophila accessory gland (Rambur et al., 2020). Thanks to its versatility as a genetic model, drosophila has been successfully used in cancer biology to decipher fundamental mechanisms in brain cancer development (Read, 2011; Read et al., 2009) as well as epithelial cancers such as lung (Levine and Cagan, 2016) and colon cancers (Bangi et al., 2016). It has even recently been used as a platform for precision medicine in cancer therapy, demonstrating its direct relevance to patient care (Bangi et al., 2019). Most prostate adenocarcinomas are thought to originate from luminal cells (Humphrey, 2012). The drosophila accessory gland is a functional equivalent of the prostate (Leiblich et al., 2012; Wilson et al., 2017; Xue and Noll, 2002) mainly composed of a monolayer of seminal fluid-secreting cells (Taniguchi et al., 2018; Xue and Noll, 2002) surrounded by a monolayer of stroma-like muscle cells encased in a strong basement membrane (Rambur et al., 2020). It is an established model to study mechanisms of epithelial prostate tumorigenesis (Ito et al., 2014; Molano-Fernández et al., 2022; Rambur et al., 2021, 2020; Vialat et al., 2024; Wilson et al., 2017). Furthermore, accessory gland differentiation and activity are driven by the drosophila sex steroid Ecdysone Receptor (EcR) (Sekar et al., 2023; Sharma et al., 2017), paralleling the role of human Androgen Receptor (AR) in the prostate physiology.

In this study, we have first interrogated prostate cancer patient expression data. We find that in Castration Resistant Prostate Cancer (CRPC), there is no evidence of canonical AR signaling reactivation; in fact, canonical AR signaling is significantly repressed in CRPC compared to primary tumors. Furthermore, cancer progression tightly correlates with this loss of canonical AR signaling. These data raise the possibility that this loss could participate to tumor resistance. However, due to the existence of non-canonical pathways, of multiple escape mechanisms and of frequent crosstalk with other steroid signaling such as Glucocorticoid Receptor, which in resistance expresses AR target genes (Arora et al., 2013), it appears difficult to study the specific effect of sex steroid canonical pathway repression on cancer progression. To study this effect, we turned to drosophila where only one steroid is known, ecdysone. So, we reproduced endocrine therapy in the drosophila accessory gland by genetically inducing tumor-specific ecdysone signaling downregulation. We show that repression of canonical sex steroid signaling actively reprograms pre-existent tumor cells to create a new tumor cell population that escapes this sex steroid deprivation. This deprivation induces a newly uncovered kind of basal extrusion, *de facto* creating a new niche for tumor development in what we define as the intrabasal compartment. This change notably relies on downregulation of EcR target gene *βTub60D*, which encodes β3 Tubulin (β3Tub). These findings possibly shed new light on the mechanisms of tumor escape.

## Results

### Canonical Androgen signaling decreases with cancer progression

#### Canonical Androgen signaling genes are downregulated in CRPC compared with primary cancer

Even though it is commonly accepted that CRPC corresponds to a phase of reactivation of AR signaling, we first assessed how much activity of canonical androgen signaling can be detected depending on the stage of prostate cancer. Because AR is a transcription factor, transcription of target genes was used as a readout across more than 1000 tumor samples from several public datasets (Abida et al., 2019; Beltran et al., 2016; Cancer Genome Atlas Research Network et al., 2015; Kumar et al., 2016; Labrecque et al., 2019; Lapuk et al., 2012; Sharp et al., 2019; Stelloo et al., 2018; Suntsova et al., 2019; “The Genotype-Tissue Expression (GTEx) project.,” 2013) and grouped in the Prostate Cancer Atlas (prostatecanceratlas.org). This tool was initially introduced by Bolis et al (Bolis et al., 2021). We first observed that *AR* expression significantly increases in primary prostate cancer compared to normal tissue, and increases again in CRPC samples compared to primary cancer (Fig. 1A). We then explored the androgen response using expression of several target genes of AR signaling (*ABCC4*, *ALDH1A3*, *AMACR*, *CAMK2*, *FASN*, *KLK2*, *KLK3*, *NKX3-1*, *PART1*, *PCA3*, *PLPP1*, *PMEPA1*, *STEAP1*, *STEAP2*, *STEAP4*). All these genes exhibit the same expression profile. Compared to normal samples, they are all significantly overexpressed in primary samples, as is *AR*. However, in CRPC samples and even though *AR* expression is still increased there, they are all significantly repressed compared to primary samples (Fig. 1B, Table 1). In fact, six of these genes (*ABCCC4*, *ALDH1A3*, *KLK2*, *KLK3*, *PART1*, *STEAP4*) are expressed at even lower levels in CRPC than in normal tissues (Fig. 1B,Table 1).

**Fig. 1.**
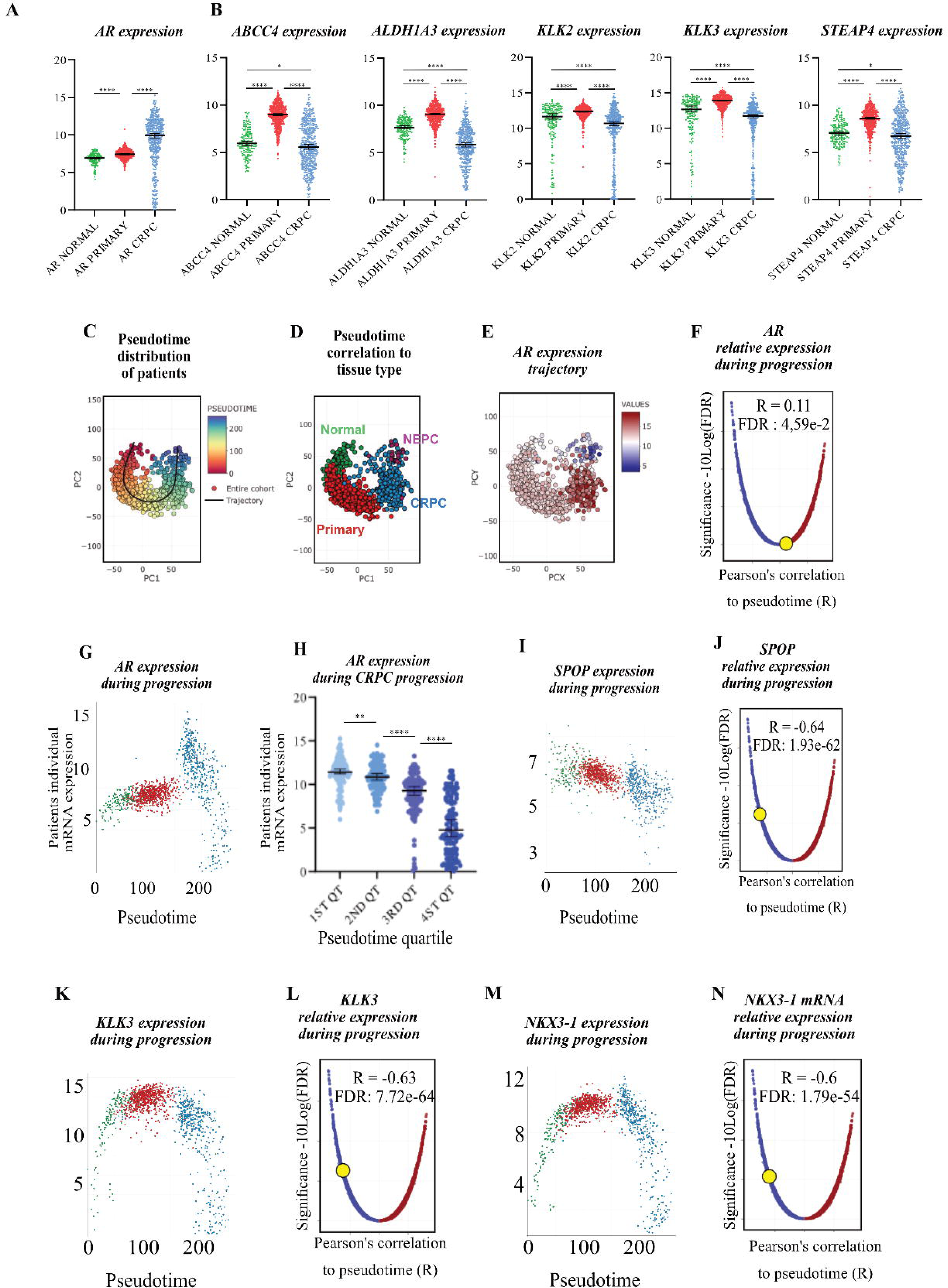
Androgen signaling activity negatively correlates with prostate cancer progression. (A-B) mRNA expression in normal (green), primary (red) or castration-resistant (CRPC) prostate cancer samples. (A) *AR* expression. (B) *AR* target genes expression. (C) Principal component analysis of sample expression defines a trajectory of disease progression with an associated pseudotime score (extracted from Prostate Cancer Atlas). (D) Pseudotime correlation segregates normal, primary, CRPC and neuroendocrine (NEPC) samples. (E) Individual *AR* expression in the samples along the trajectory. (F) Relative change of expression of AR during progression. X-axis represents Pearson’s correlation coefficient between mRNAs and pseudotime as Y-axis displays the associated significance adjusted for false discovery rate (FDR). (G) Individual *AR* expression in the samples depending on pseudotime (X-axis) and sample type (color code). (H) CRPC samples are separated in four quartiles (1ST to 4TH QT) depending on their pseudotime. *AR* expression is then plotted for individual samples. (I-N) Individual expression and relative change of expression of three AR target genes (SPOP, KLK3, NKX3-1) during progression. * p<0.05; ** p<0.01; ****p<0.0001.

**Table 1.**
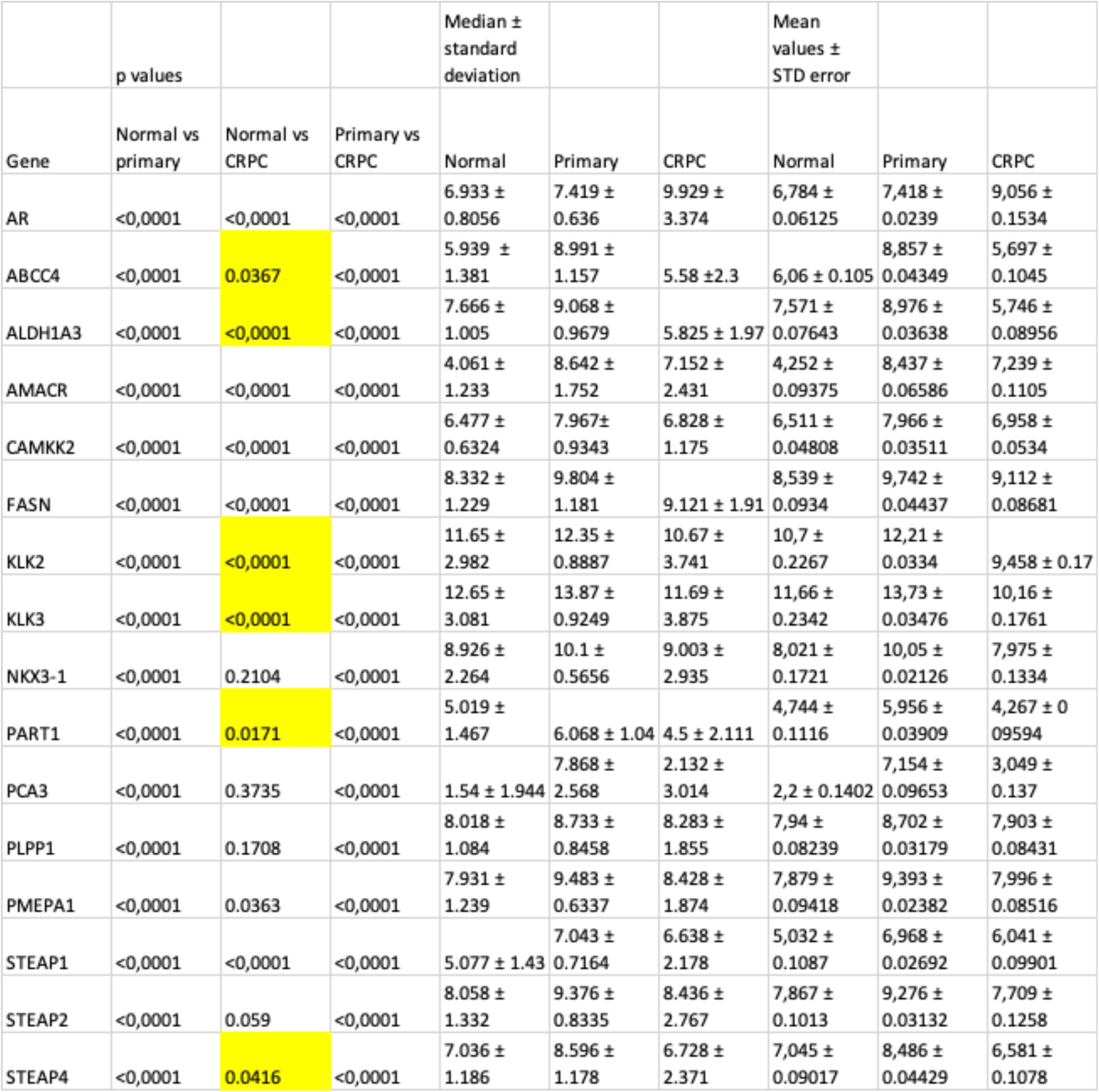
Canonical AR Signaling genes are downregulated in CRPC compared with primary cancer. Based on 173 Normal samples, 708 Primary cancer samples and 484 CRPC samples. In yellow: six of the genes are significantly less expressed in CRPC than in normal samples.

#### Canonical Androgen signaling gene expression negatively correlates with prostate cancer progression

The Prostate Cancer Atlas also correlates changes in transcriptomic profiles with cancer progression (i.e. pseudotime progression, Fig.1C), with a high degree of confidence, allowing retrospective association of specific changes in gene expression with the cancer trajectory.

Logically, plotting individual patients on this trajectory discriminates normal tissue (green, low pseudotime, Median ± Std error = 54.58 ± 2.396), primary cancer (red, intermediate pseudotime = 105.9 ± 0.7952) and CRPC (blue, high pseudotime = 196.5 ± 1.322) among which the most aggressive form -neuroendocrine cancers (NEPC, violet)- clusters at the final tip of the trajectory (Fig. 1D). *AR* expression itself peaks in CRPC samples (Fig. 1E) but only weakly correlates with progression (Fig.1F, R = 0.11, FDR = 0.0459). To understand why this correlation is so weak, we plotted *AR* expression against pseudotime (Fig. 1G). *AR* expression there appears to vary widely in CRPC samples. We therefore subdivided CRPC samples into four groups based on pseudotime, i.e., their state of progression (Fig. 1H). The 1st quartile (1ST QT) displays mean AR expression of 11.41 ± 0.16; 2ND QT is significantly lower at 10.83 ± 0.15 (p = 0.0209). Decrease from quartile to quartile continues for 3RD QT (8.82 ± 0.22, p<0.0001) and 4TH QT (5.27 ± 0.31, p<0.0001). Thus, even though *AR* expression clearly increases in primary cancer and then CRPC, its expression steadily decreases within CRPC when state of progression is considered.

At the same time, canonical androgen signaling appears as the most downregulated pathway along cancer progression -especially in CRPC samples (Bolis et al., 2021)- as shown for *SPOP*, one of the most frequently mutated genes in prostate cancer and a key player in genome integrity (Barbieri et al., 2012; Boysen et al., 2015) (Fig. 1I-J, R = -0.64, p = 1.93e-52), *KLK3* -coding for the Prostate Specific Antigen (Fig. 1K-L, R = -0.63, p = 7.72e-64)- or *NKX3-1,* which is itself a specific prostate tumor suppressor gene (Bhatia-Gaur et al., 1999) (Fig. 1M-N, R = -0.60, p = 1.79e-54). Overall, even though *AR* itself is globally more expressed in CRPC patients, canonical androgen signaling definitely decreases with cancer progression.

#### CRPC-specific loss of correlation between AR expression and canonical AR signaling

To understand this phenomenon, we plotted expression of *AR* expression against its target genes (here *ALDH1A3*) for normal, primary cancer or CRPC samples (Fig. 2A). In normal tissues, *AR* and *ALDH1A3* expression are concentrated in a relatively narrow range, making all samples look alike (Fig. 2B). In primary cancer, the increase in *AR* expression (from 6.93 ± 0.81 to 7.42 ± 0.64, p<0.0001) correlates with an increase in *ALDH1A3* expression (from 7.67 ± 1 to 9.07 ± 0.97, p<0.0001), keeping samples close to each other but shifted to a higher expression range compared to normal samples (Fig. 2C). However, in CRPC samples, despite a still higher *AR* median expression (from 7.42 ± 0.64 to 9.93 ± 3.37, p<0.0001), individual *AR* expression is dispersed over more than 10 logs (from 0.11 to 13.16). *ALDH1A3* expression also varies hugely across samples, but with a Pearson Coefficient of 0.494, remains weakly correlated with AR expression in individual samples. However, *ALDH1A3* expression does strongly decrease (from 9.07 ± 0.97 to 5.83 ± 1.97, p<0.0001) in CRPC (Fig. 2D), indicating that AR does not appear to activate expression of this gene as efficiently as in normal or primary cancer tissues.

**Fig. 2.**
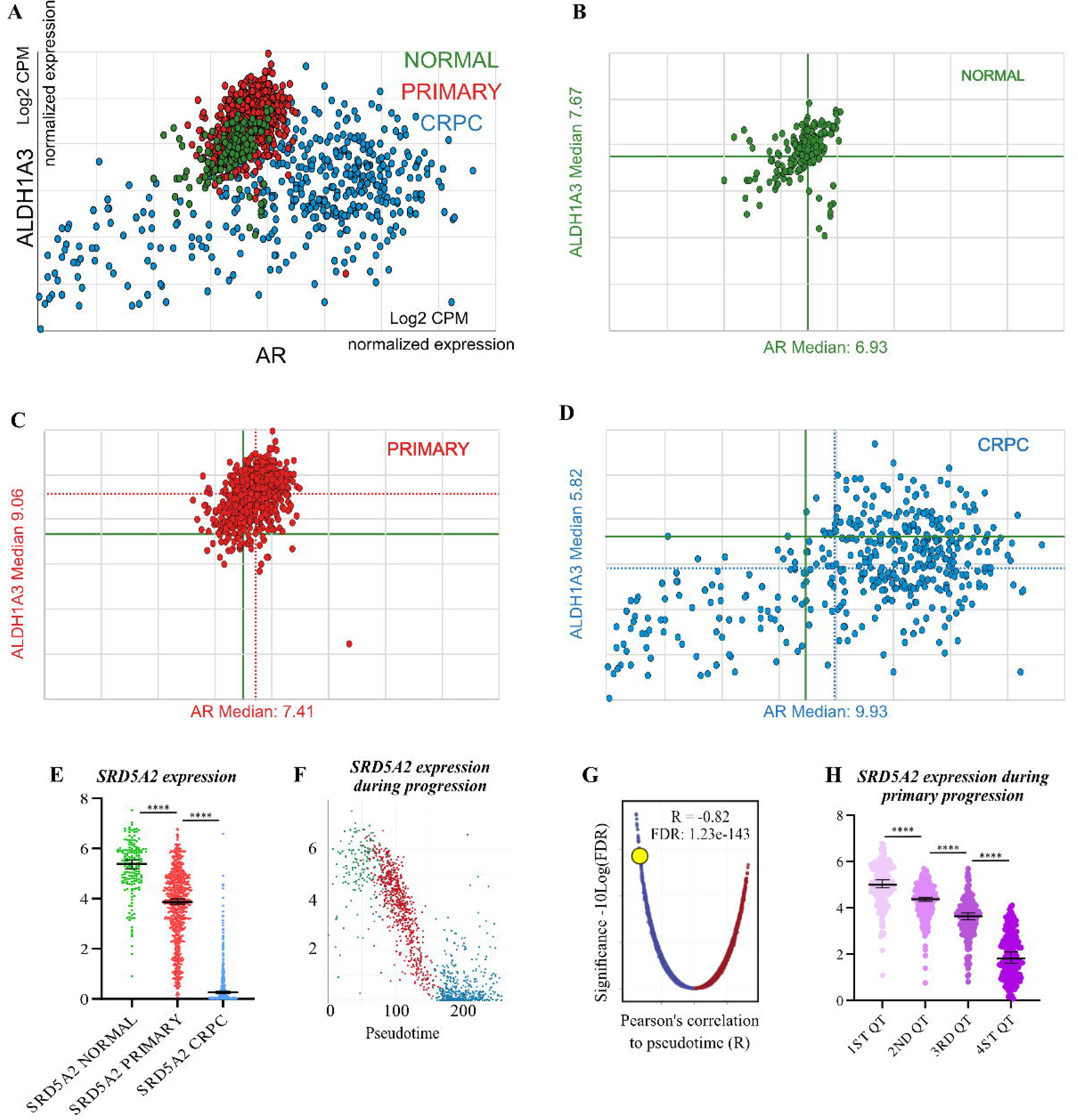
Loss of correlation between AR expression and canonical AR signaling coincides with downregulated expression of *SRD5A2*. (A) Individual *AR* expression (X-axis) versus ALDHIA3 expression (Y-axis) in the samples depending on sample type (color code). (B-D) Same expression separated by sample type. (E) *SRD5A2* -coding for the DHT-producing enzyme-mRNA expression in normal (green), primary (red) or castration-resistant (CRPC) prostate cancer samples. (F-G) Individual expression and relative change of expression of *SRD5A2* during progression. (H) Primary samples are separated in four quartiles (1ST to 4TH QT) depending on their pseudotime. *SRD5A2* expression is then plotted for individual samples. ****p<0.0001.

We then wondered whether androgen metabolism itself could be disturbed. Interestingly, *SRD5A2*, which encodes the enzyme responsible for the production of the active metabolite of testosterone, DiHydroTestosterone (DHT) (Andersson et al., 1991), is downregulated in primary samples and even more so in CRPC samples (Fig. 2E). *SRD5A2* is in fact one the most downregulated genes during cancer progression (Fig. 2F, Fig. 2G, R=-0,82, p=1.23e-143). However, its progressive loss of expression starts from primary cancer (Fig. 2H).

Potentially, the lack of response to the peak of AR expression in early CRPC could result from reduced DHT metabolization that begins in primary tumors. Indeed, DHT levels are lower in untreated cancer tissues than in benign prostatic hyperplasia (Geller et al., 1978). Furthermore, in men suspected of prostate cancer who underwent needle biopsies, a high percentage of cancer-positive cores -associated with poor prognosis- also correlates with lower DHT levels (Miyoshi et al., 2014). Thus, DHT concentration decreases in prostate cancer even before castration and castration resistance, and correlates with poor prognosis. Overall, transcript analysis of major prostate cancer cohorts indicates that canonical Androgen Receptor signaling negatively correlates with cancer progression, even in samples classified as hormone refractory in which androgen signaling is generally considered active.

### Tumor escape induced by Ecdysone Receptor downregulation

Considering these results, we asked whether repression of canonical sex steroid signaling could by itself induce cancer progression. To test this hypothesis, we turned to a drosophila model of prostate cancer we previously engineered, in which we can also study in detail the phenomenon of basal extrusion, considered as the founding step of cancer formation.

#### Epithelial tumor cells and extra-epithelial tumors in EGFR*λ*-mediated tumorigenesis

Thanks to genetic tools available in drosophila based on the UAS/Gal4 expression system, it is possible to reproduce tumor initiation in the drosophila accessory gland. For this, clonal expression of a constitutively active form of EGFR -EGFR*λ*- is realized in ∼1% of the main cells of the accessory gland. This oncogene is co-expressed with GFP, allowing tracing of tumor cells. To provide an adequate control for tumorigenesis (EGFR*λ* CTL), it is furthermore co-expressed with a second nuclear GFP, as every condition must bear the same number of UAS elements to avoid Gal4 titration effect. Under these conditions, tumor initiation displays different outcomes. Some clonal cells undergo proliferation at low rates but do not migrate, and thus remain inside the epithelium (Fig. S1A-F, *i.e.* clones, white arrowheads). Among these cells, some display disorganized cell-cell interaction markers, indicating disturbed epithelial differentiation and probable loss of adhesion with neighbor cells (Fig. S1A-C, thick empty arrowheads). Due to their mild hyperproliferation, these cells can be considered part of benign tumors. Finally, some clonal cells migrate and develop into extra-epithelial tumors (Fig. S1D-F, empty arrowheads), where cells are completely devoid of epithelial markers (Fig. S1F), indicating in drosophila a phenomenon of Epithelial-Mesenchymal Transition (EMT).

As demonstrated by their presence outside the muscle sheet that encloses the gland (revealed by F-actin staining in Fig. 3A and later), these tumor cells invade through the basement membrane, a behavior known as epithelial basal extrusion that is specific to cancer cells according to the National Cancer Institute. We previously showed that these extra-epithelial tumors harbor phenotypic abnormalities typically associated with tumor development, such as hyperplasia, hypertrophy, loss of epithelial marker expression (Fig. S1F, Fig. 3) and promotion of angiogenesis-like tracheogenesis. We also previously described that many of them retain a specific feature of accessory gland epithelial cells -twin nuclei-confirming that these cells originate from the accessory gland (Rambur et al., 2020).

**Fig. 3.**
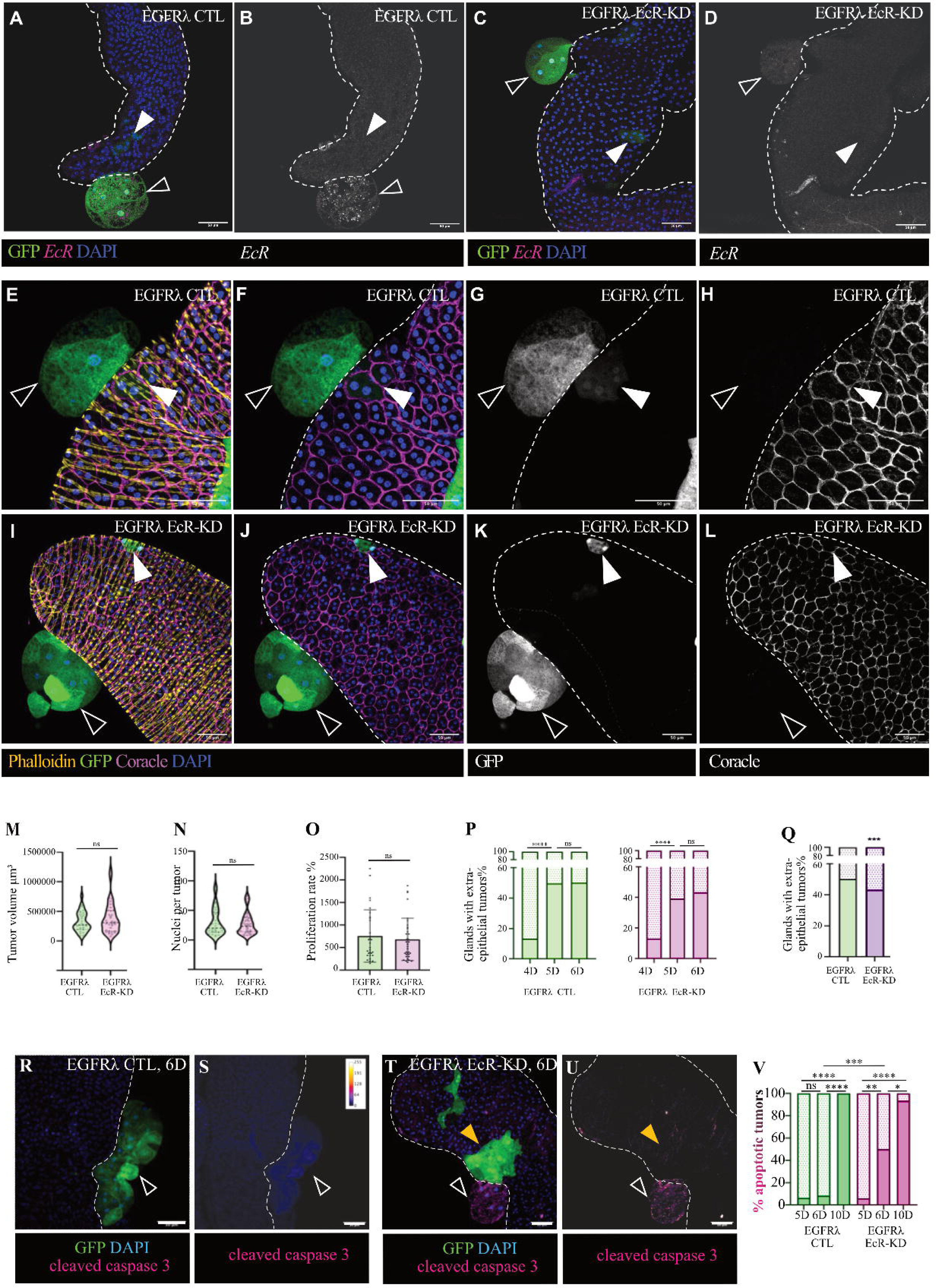
Treatment-like effect induced by Ecdysone Receptor downregulation. (A-L and R-U) Confocal imaging of accessory glands (delimited by white dashes). White arrowheads indicate epithelial clones. Empty arrowheads indicate extra-epithelial tumors. Yellow arrowheads indicate a tumor cell population specific to EGFR*λ* EcR-KD genotype. (A-D) FISH detection of EcR mRNA in EGFR*λ* CTL and EGFR*λ* EcR-KD genotypes. (E-L) Shared phenotypes between the two genotypes. (M-O) Extra-epithelial tumor characteristics defined by 3D-reconstruction. (P) % of glands bearing extra-epithelial tumors 4, 5 or 6 days after clonal induction for the different genotypes. (Q) Comparison of the maximal extra-epithelial tumors percentage (at 6D) between the two genotypes. (R-U) Cleaved caspase 3 staining in the different tumor cell populations. (V) Quantification of the number of apoptotic extra-epithelial tumors at 5, 6 or 10 days after clonal induction for the different genotypes. Scale bars: 50μm. Number of experiments (N) and of individual flies (n) are given in Table 2. Statistical tests are explained in the Methods section; ns: non-significant; * p<0.05; ** p<0.01; *** p<0.001; ****p<0.0001.

#### Treatment-like effect induced by Ecdysone Receptor downregulation

We further evaluated *EcR* expression during tumorigenesis. By RNA FISH, we show that *EcR* is overexpressed in tumor cells that were able to extrude basally (Fig. 3A-B). This phenocopies *AR* overexpression in human prostate cancer and raises questions about the potential role of EcR in accessory gland tumorigenesis. We so co-expressed an RNAi targeting EcR with the oncogene (EGFR*λ* EcR-KD, Fig. 3C-D). At first sight, this co-expression induces growth of epithelial clones and extra-epithelial tumors similar to the control (compare white and empty arrowheads in Fig. 3E-H to Fig. 3I-L). At 6D, tridimensional reconstruction of image stacks indicates that the tumors display the same volume and the same number of cells in both conditions (Fig. 3M, N). Thus, these tumors display similar proliferation rate of ∼700% compared with clones expressing GFP only (Fig. 3O), which are typically composed of two cells at six days of induction (Fig. S3A). We conclude that phenotypic development of extra-epithelial tumors does not rely on ecdysone signaling. We then examined whether extra-epithelial tumor formation itself is modified.

Temporal dynamics of extra-epithelial tumor formation is not affected by EcR RNAi co-expression (Fig. 3P): extra-epithelial tumors typically start to appear between the third and fourth days after clonal induction (3D and 4D), peaking at 5D regardless of genotype, indicating that basal extrusion is active only during this short period of time. However, a modest but significant decrease in the total number of extra-epithelial tumors is detected upon EcR downregulation (Fig. 3Q). Together, these data show that loss of ecdysone signaling specifically affects the overall quantity of tumor basal extrusion, but has neither an effect on its temporal dynamics nor on tumor growth rate outside the gland.

In human, sex steroid deprivation also induces death of treatment-sensitive tumor cells (Dowsett et al., 2006; Isaacs et al., 1994). 6 days after clone induction (6D), cleaved-caspase 3 staining reveals that, in the control condition, tumor cells do not present signs of apoptosis (Fig. 3R-S). However, at the same time point, EcR RNAi co-expression significantly induces apoptosis in extra-epithelial tumors (Fig. 3T-U, empty arrowheads in left panels; middle columns in Fig. 3V for quantification). This indicates that, in the drosophila accessory gland, some tumor cells rely on sex steroid signaling for their survival, as in the human disease.

Interestingly, at 10 days after induction (10D), most extra-epithelial tumors harbor apoptotic staining, independently of genotype (Fig. 3V, right columns). Thus, despite their ability to undergo the complex program of basal extrusion, these cells seem unable to survive for long periods, indicating limited tumor potential.

Together, these results indicate that repression of EcR in accessory gland tumors cells displays a treatment-like effect: extra-epithelial tumor formation is (slightly but) significantly impaired, and cells composing these tumors are sensitive to this deprivation and undergo apoptosis.

#### Tumor escape induced by Ecdysone Receptor downregulation

Strikingly, in EGFR*λ* EcR-KD condition, we also detect a new type of tumorigenesis: large tumor masses now grow inside the glands (Fig. 3T, Fig. 4A-D, yellow arrowheads). Thus, generation of this new phenotype apparently depends solely on lack of ecdysone signaling in tumor cells. Contrary to the cells that remain in the epithelium in the EGFR*λ* CTL condition, these cells generally express low levels of epithelial markers (Fig. 4D), indicating a loss of epithelial differentiation. Notably, unlike extra-epithelial tumors, this new tumor cell population shows no sign of apoptosis regardless of induction time (Fig. 4E-H), and tend to invade the whole gland, unlike clone cells in the EGFR*λ* CTL condition (white arrowheads in Fig. 4E-F, yellow arrowheads in Fig. 4G-H). Thus, this new population displays an increased proliferative capacity compared with the EGFR*λ* CTL condition. Together, this indicates that these cells are not only escaping EcR repression but also now thrive over time under this stress.

**Fig. 4.**
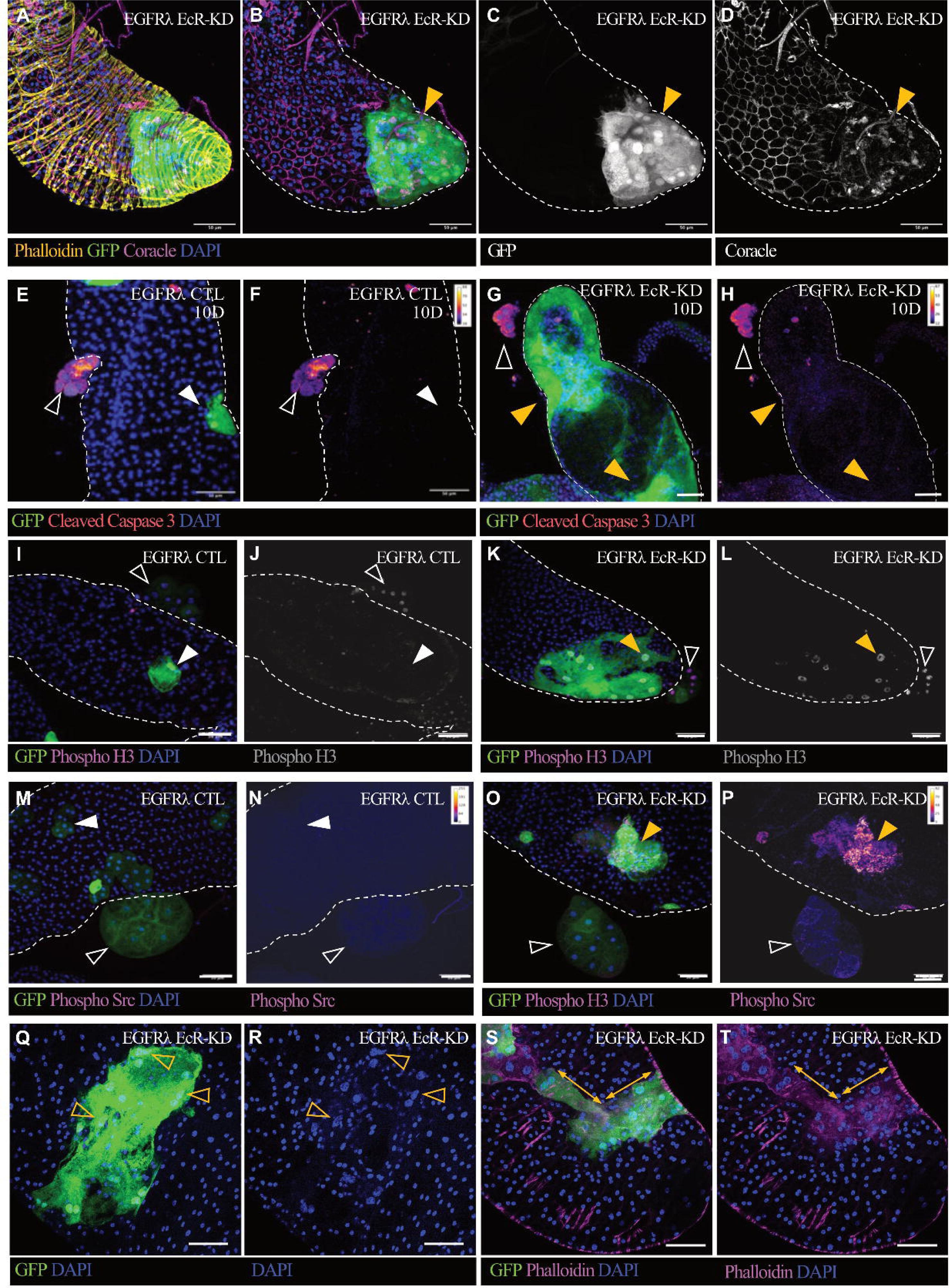
Tumor escape induced by Ecdysone Receptor downregulation. (A-T) Confocal imaging of accessory glands (delimited by white dashes in some images). White arrowheads indicate epithelial clones. Empty arrowheads indicate extra-epithelial tumors. Yellow arrowheads indicate the new tumor cell population which is specific to EGFR*λ* EcR-KD genotype. (A-D) Coracle staining indicates epithelial status of the cells. (E-H) Cleaved caspase 3 staining in the different tumor cell populations 10 days after clonal induction for the different genotypes. (I-L) Phospho-Histone 3 staining 6 days after clonal induction for the different genotypes. (M-P) Phospho-Src staining 6 days after clonal induction for the different genotypes. (Q-R) DAPI staining reveals nuclear shapes of cells of the new population (yellow empty arrowheads). (S-T) Multiple axes of growth (double-headed arrows) and F-actin reorganization (phalloidin staining, magenta) in the new tumor cell population. Scale bars: 50μm.

#### Markers of higher progression of the new tumor cell population

Contrary to tumor cells in EGFR*λ* CTL condition, this new cell population displays strong staining for Ser10 Phospho-Histone 3 (pH3), a marker of late phases of the cell cycle prior to cell division (compare white to yellow arrowheads in Fig. 4G-H to 4I-J). Furthermore, they exhibit a positive staining for phosphorylated proto-oncogene tyrosine-protein kinase Src (pSrc), a classical marker of aggressiveness (compare white to yellow arrowheads in Fig. 4K-L to 4M-N) (Baumgartner et al., 2008). These tumor cells also display many typical defects, such as a clear pleomorphism with either compact or fusiform cells, and strongly disorganized clones (Fig. 4Q-T). In the same clone and even within the same cell, nuclei differ markedly in size (anisocaryosis) with frequent hypertrophied nuclei (macrocaryosis). Nuclei shapes are often altered (empty arrowheads in Fig. 4Q-R). Clones can grow along multiple axes (double-headed arrowheads in Fig. 4S-T) and present a strong fibrillar framework (Fig. 4T). Together, these results highlight that this new population has reached a higher step of tumor progression compared to previously described clones and extra-epithelial tumors, which form rounder, well-delimited masses.

### Basal extrusion is modified in case of tumor escape

We then asked how these new tumor cells appear inside the glands.

#### A second kind of basal extrusion unveiled by EcR downregulation

In EGFR*λ* CTL condition (Fig. 5A-D, white arrowheads), as in case of co-repression of Ras/MAPK, PI3K/Akt (Rambur et al., 2020) or cholesterol metabolic pathway (Vialat et al., 2024), tumor clones that appear in the gland remain strictly in the epithelium (orthogonal view in Fig. 5C-D, where phalloidin staining (yellow) indicates the position of the muscle layer within the basement membrane). By contrast, in EGFR*λ* EcR-KD condition, the new population of tumor cells grows between the normal epithelium (delineated by white dashes in the orthogonal views in Fig. 5G-H) and the basement membrane (Fig. 5E-H, yellow arrowheads). Thus, these cells were able to migrate basally out of the normal epithelium, thereby pushing it into the gland lumen. At the same time, they did not cross the basement membrane. This redefines basal extrusion as two different phenomena with a common start. In this first step, cells extrude basally from the epithelium. Then, cells can either cross the basement membrane and form extra-epithelial tumors or, depending on *EcR* repression, grow inside the gland. Blocking ecdysone signaling therefore promotes a new tumor cell fate, in which tumor cells accumulate between the normal epithelium and the basement membrane, forming highly proliferative tumors that escape steroid signaling downregulation in what we define as the intrabasal (IB) compartment.

**Fig. 5.**
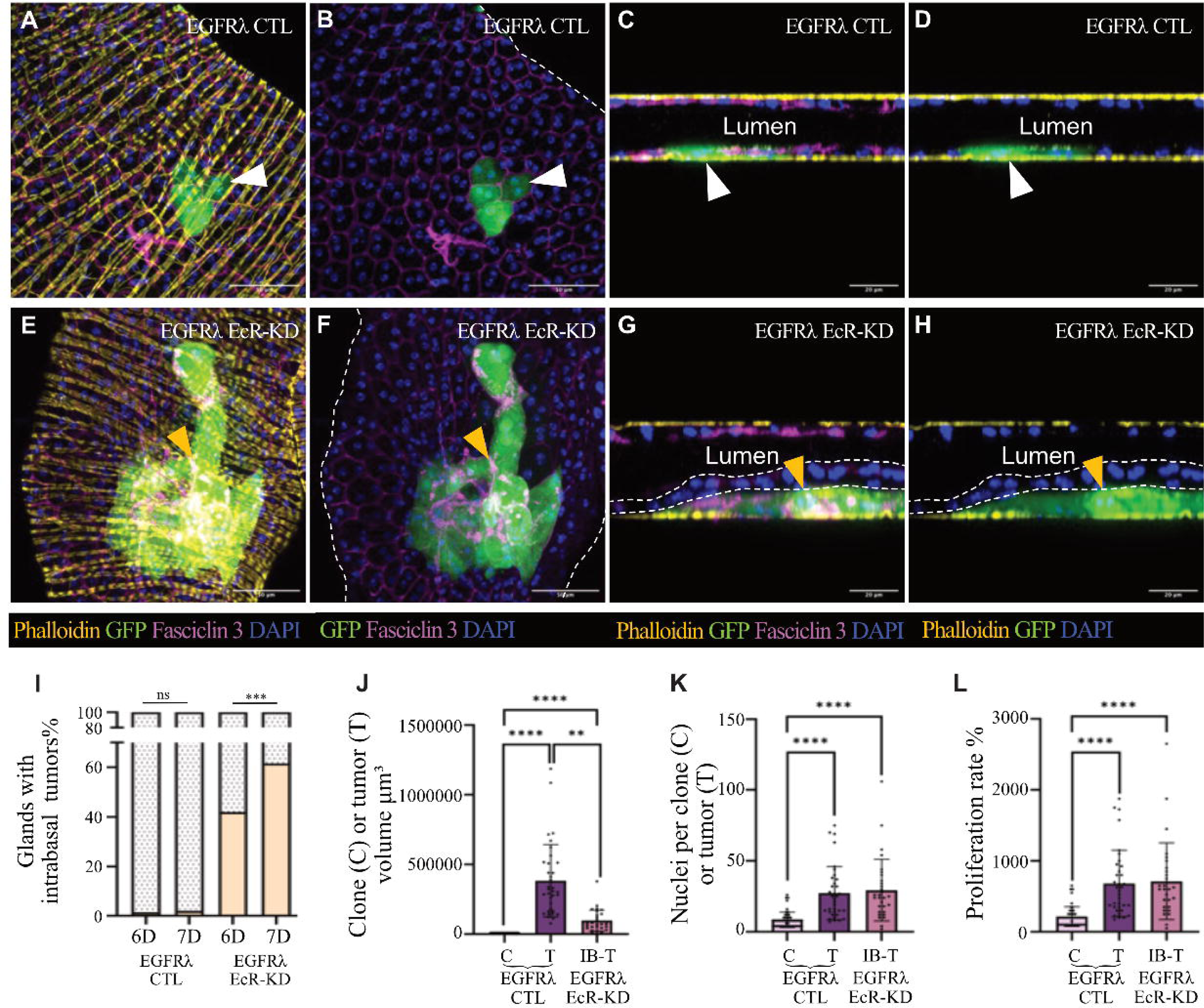
Basal extrusion is modified in case of tumor escape. (A-T) Confocal imaging of accessory glands. White arrowheads indicate epithelial clones. Empty arrowheads indicate extra-epithelial tumors. Yellow arrowheads indicate the new tumor cell population which is specific to EGFR*λ* EcR-KD genotype. (A-B, E-F) XYZ imaging. Glands are delimited by white dashes in B and F. (C-D, G-H) XZY orthogonal reconstructions of the glands, showing the lumens and in-depth view of the epithelium. Epithelium pushed into the lumen is delimited by white dashes in G and H, as intrabasal tumor cells lay between this epithelium and the basement membrane (yellow). (I) % of glands bearing intrabasal tumors 6 or 7 days after clonal induction for the different genotypes. (J-L) Comparison of intrabasal vs other tumor characteristics defined by 3D-reconstruction (C: clones; T: extra-epithelial tumors; IB-T: intrabasal tumors). Scale bars: 50μm in XYZ views and 20μm in XZY reconstructions. ns: non-significant; ** p<0.01; *** p<0.001; ****p<0.0001.

#### Intrabasal growth relies on a new, inducible kind of basal extrusion

Given that we concomitantly detected a slight decrease in extra-epithelial tumor formation in the EGFR*λ* EcR-KD condition (Fig. 3Q), we asked whether formation of intrabasal tumors arises from a redirection from full basal extrusion to partial basal extrusion. We show that, in contrast to extra-epithelial tumors, intrabasal tumors continue to appear and grow over time, as more glands display at least one intrabasal tumor at 7D than at 6D (Fig. 5I); thus, they apparently do not replace lost extra-epithelial tumors. As a result of this growth, at 7D there are already significantly more glands displaying an intrabasal tumor that the maximal percentage of extra-epithelial tumors achieved in the EGFR*λ* CTL genotype (61.6% vs 49.6%, p<0.001. N=29, n=1117 vs N=3, n=268). This shows that, in EGFR*λ* EcR-KD condition, formation of intrabasal tumors does not result from mere inhibition of full basal extrusion but from promotion of a different kind of basal extrusion that ultimately generates tumor escape.

We then characterized these intrabasal tumors. At 6D, intrabasal tumors (IB-T) are significantly smaller than extra-epithelial tumors (T) but still significantly larger than epithelial clones of EGFR*λ* CTL genotype (C) (Fig. 5J). This increase in volume mostly results from an increased proliferation rate and, consequently, increased tumor cell number in intrabasal tumors, which at 6D already compares with that of extra-epithelial tumors (Fig. 5K-L). Thus, considering the total percentages of extra-epithelial and intrabasal tumors, *EcR* downregulation more than doubled the number of tumor cells during the course of the experiment.

Overall, *EcR* knockdown in tumor cells displays two opposite effects. On the one hand, it induces apoptosis of extra-epithelial tumors and modestly decreases their rate of formation. On the other hand, it strongly induces a new kind of tumorigenesis, which develops and thrives in the newly uncovered intrabasal compartment.

### Ecdysone-dependent molecular control of tumor escape and basal extrusion

We then sought to determine the molecular mechanisms linking EcR Signaling to tumorigenesis. Prostate tumor escape is generally linked to resistance to androgen deprivation where AR is able to be activated despite hormone deprivation, through activating mutations in the gene or ectopic sex steroid production by tumors or their surrounding tissues. However, patients data indicate a decreased canonical androgen signaling in CRPC (Fig. 1-2). It is therefore important to understand whether the observed escape results from a mechanism of resistance or from a genuine lack of ecdysone signaling. We therefore downregulated multiple targets at different levels of this pathway in a normal or tumoral context (Fig. S2).

#### Loss of ecdysone signaling does not promote tumorigenesis by itself

First, we tested the effect of a loss of ecdysone signaling in a non-tumoral context. As previously shown (Rambur et al., 2020), at 6D, clones expressing GFP are composed of two cells (Fig. S3A). Downregulation of EcR expression (EcR RNAi) or impairment of ecdysone synthesis (Shadow RNAi, Sad-KD, see Fig. S2) induces the same effect: as in CTL, clones are composed of two cells that appear of normal shape and nuclear number (Compare Fig. S3A and S1B and S3C). No tumorigenesis is detected (Fig. S3D). The only detected phenotype is a slight decrease in epithelial cell growth without change in proliferation (Fig. S3E-G). This hypotrophy could reflect a partial loss of cells’ ability to produce seminal fluid, which relies on ecdysone signaling in accessory gland cells.

#### Lack of canonical ecdysone signaling drives tumor escape

Paralleling human metabolization of testosterone into active DiHydroTestosterone (DHT) by the SRD5A2 gene product, ecdysone must be metabolized into active 20-hydroxyecdysone by Shade (Shd) (Petryk et al., 2003). As *SRD5A2* dramatically decreases during cancer progression (Fig. 2E-H), we downregulated *Shd* in tumor cells. Tumor progression is phenocopied in glands of the EGFR*λ* Shd-KD genotype compared with the EGFR*λ* EcR-KD condition, for both the decrease in extra-epithelial tumors and the growth of intrabasal tumors (Fig. 6A, magenta columns). By targeting an independent gene, this confirms that lack of EcR signaling leads to tumor resistance in the drosophila accessory gland.

**Fig. 6.**
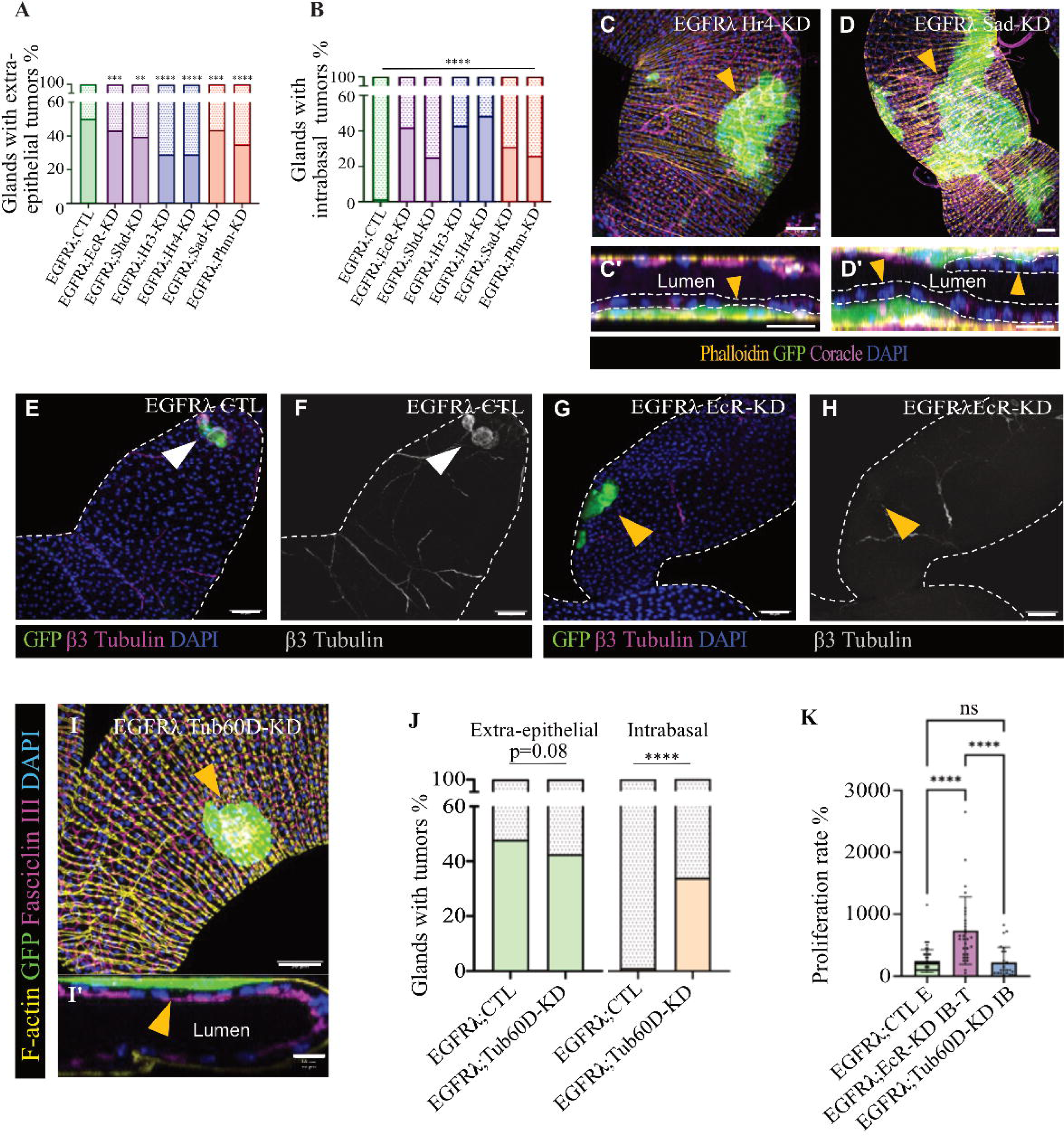
Ecdysone-dependent molecular control of tumor escape and basal extrusion. (A-B) % of glands bearing extra-epithelial (A) or intrabasal tumors (B) 6 days after clonal induction for different genotypes. (C-D’) Confocal imaging of accessory glands. Yellow arrowheads indicate intrabasal tumors. (C, D) XYZ imaging. (C’, D’) Corresponding XZY orthogonal reconstructions of the glands, showing the lumens and in-depth view of the epithelium. Epithelium pushed into the lumen is delimited by white dashes. (E-H) β3 Tubulin staining 6 days after clonal induction for different genotypes. Glands are delimited by white dashes, white arrowheads indicate tumors clones. (I-I’) Confocal imaging of accessory glands. (I) XYZ imaging. (I’) Corresponding XZY orthogonal reconstruction of the glands. (J) % of glands bearing extra-epithelial (left) or intrabasal tumors (right) 6 days after clonal induction for different genotypes. (K) Comparison of proliferation rates defined by 3D-reconstruction (E: epithelial clones; IB-T: intrabasal tumors; IB: intrabasal cells in EGFR*λ* Tub60D-KD genotype). Scale bars: 50μm in XYZ views and 20μm in XZY reconstructions. ** p<0.01; *** p<0.001; ****p<0.0001.

Furthermore, ecdysone signaling relies on recruitment of specific EcR target genes -the early-late transcription factors Hr3 and Hr4- to drive ecdysone responses (Carney et al., 1997; Gauhar et al., 2009). Again, tumor progression in both EGFR*λ* Hr3-KD and EGFR*λ* Hr4-KD conditions is strikingly similar to the EGFR*λ* EcR-KD condition (Fig. 6A-B, blue columns, Fig. 6C-C’, Fig. S3), showing that lack of canonical ecdysone signaling leads to tumor escape.

#### Complete dependence on tumor-produced Ecdysone to alleviate tumor escape

Prostate cancer cells are able to produce steroids (Locke et al., 2008), which contribute to AR reactivation in castration-resistant prostate cancer (CRPC) (Cai et al., 2011). In drosophila, the Halloween genes encode the enzymes that synthetize ecdysone. Phantom (Phm) is the limiting enzyme for this synthesis, whereas Shadow (Sad) is the terminal enzyme producing ecdysone. The EGFR*λ* Phm-KD and EGFR*λ* Sad-KD conditions again phenocopy the EGFR*λ* EcR-KD condition (Fig. 6A-B, red columns, Fig. 6D-D’, Fig. S4). This shows -by targeting still two other independent genes and still another process- that, in the drosophila accessory gland, growth of intrabasal tumors results from lack of ecdysone signaling.

Furthermore, this demonstrates that circulating ecdysone cannot compensate for lack of tumor-produced ecdysone, which is therefore the only indispensable resource to prevent tumor escape in the accessory gland. We previously demonstrated that autocrine loops recruiting Ras/MAPK and PI3K/Akt pathways are established to support basal extrusion (Rambur et al., 2020). We show here that an autocrine/intracrine loop is also established to activate ecdysone signaling and sustain the same phenomenon. However, blocking this loop now also induces growth of intrabasal tumors, in contrast to inhibition of Ras/MAPK and PI3K/Akt pathways. Thus, promotion of this mechanism of tumor escape is specifically induced by the loss of ecdysone signaling in these cells.

#### EcR target gene Tub60D specifically represses intrabasal growth

We then investigated molecular drivers that could promote intrabasal growth. Cytoskeletal reorganization is a major determinant of tumor cell invasion and resistance, and intrabasal tumor cells harbor profound cytoskeletal change. *Beta Tubulin at 60D* (*Tub60D*) encodes β3 Tubulin and is a target of ecdysone signaling (Bruhat et al., 1990). β3 Tubulin accumulates in intra-epithelial clones of EGFR*λ* CTL condition but is mostly absent in intrabasal tumor cells (compare Fig. 6E-F to G-H). Strikingly, downregulation of *Tub60D* (EGFR*λ* Tub60D-KD condition) is sufficient to induce intrabasal growth (Fig. 6I-I’, Fig. 6J, right columns), but overall has no effect on formation of extra-epithelial tumors (Fig. 6J, left columns).

Furthermore, these extra-epithelial tumors remain unchanged compared to the other tested conditions (Fig. S6). Thus, in this condition as well, intrabasal growth represents an inducible program that is completely independent of extra-epithelial tumor formation.

Furthermore, in the EGFR*λ* Tub60D-KD condition, intrabasal tumors do not proliferate more than epithelial clones (Fig. 6K). Lack of β3 Tubulin therefore appears as a major determinant of intrabasal growth but does not promote any advantage by itself, even in a tumoral context. This indicates that the ability to achieve escape relies, in the drosophila accessory gland, on ecdysone signaling deprivation and not merely on the ability of tumor cells to reach the intrabasal compartment.

## Discussion

The National Cancer Institute defines cancer by their abnormal cells dividing without control and invading nearby tissues. Epithelial cancers represent ∼90% of all cancers (Bray et al., 2018). As epithelial cells rest on a strong extracellular matrix on their basal side, epithelial tumor cells must cross this basement membrane to invade nearby tissues. This is why this step of invasion is called basal extrusion. Little is known about basal extrusion when it occurs physiologically during development and when it occurs in the pathophysiological context of tumorigenesis. Basal extrusion appears as a complex process: it can be induced by the clonal expression of different oncogenes (Anton et al., 2018; Rambur et al., 2020; Villeneuve et al., 2019); may occur in balance with apical extrusion (Anton et al., 2018; Marshall et al., 2011; Ventrella et al., 2023); relies on multiple signaling pathways such as Notch (Ventrella et al., 2023), Ras/MAPK and PI3K/Akt (Rambur et al., 2020) or PKC activation (Villeneuve et al., 2019); depends on cholesterol uptake (Vialat et al., 2024) and lipid raft activity (Kajiwara et al., 2022); and can be promoted by extracellular matrix degradation even though, counterintuitively, this does not seem to be a necessary step (Shen et al., 2014). It also depends on profound remodeling of cytoskeletal organization and of cell-cell and cell-matrix interactions (Chountala et al., 2012; Ganier et al., 2018; Melo et al., 2023; Slattum et al., 2009; Villeneuve et al., 2019). Overall, our data indicate that basal extrusion of tumor cells appears furthermore as a two-fate phenomenon that, in drosophila accessory gland, is tightly regulated by ecdysone signaling, as downregulation of this signaling induces growth of intrabasal tumors. Interestingly, clonal expression of an oncogene in mouse epithelia induces formation of dome-like structures where tumor cells accumulate at the same localization, showing that this precise tumor fate also exists in mammals (Shirai et al., 2022). This localization has long been known to favor tumor growth, as exemplified by the widespread use of subcapsular renal xenografting in cancer research (Bogden et al., 1979). Moreover, in a mouse model of intraductal human breast carcinogenesis, Zoeller et al demonstrated that tumor cells adjacent to the basement membrane display extra-resistance to treatments when compared with others, indicating that this specific microenvironment is most conducive to survival upon treatment (Zoeller et al., 2017). An ability for tumor cells to reach this compartment could then confer them a higher chance of survival and progression.

However, we were able to push tumor cell growth in this compartment by manipulating *Tub60D* expression. In this case, no increase in cell proliferation and no apparent cell disorganization are associated with intrabasal growth. Thus, by itself, growth in the intrabasal compartment is not sufficient for tumor progression in the accessory gland.

Conversely, in this context, tumor cell resistance is actively induced by decreased canonical ecdysone signaling. And, in human, we also show that AR signaling decreases with prostate cancer progression. This lack of AR signaling also favors the most aggressive NEPC (Turpin et al., 2023). In breast cancer, ER loss is a common event largely associated with tumor aggressiveness (Kuukasjärvi et al., 1996). In prostate cancer, in the numerous alterations that can affect AR, many mutations found in tumors are loss of function (Hay and McEwan, 2012). Moreover, in human, Androgen Deprivation Therapy (ADT) shows opposite effects depending on cancer stage. For poorly differentiated or metastatic tumors, ADT has largely proven its efficacy by increasing patient survival. However, for localized prostate cancer, some studies argue that ADT displays no beneficial effect and is even associated with worse outcome than active surveillance for patient survival (Lu-Yao et al., 2008; Merglen et al., 2007). In men with hypogonadism, testosterone supplementation therapy has been feared to increase prostate cancer risk. However, reviews of the scientific literature show no increase in prostate cancer incidence with this treatment (Dupree et al., 2014; Michaud et al., 2015).

Finally, in High-Grade Prostate Intraepithelial Neoplasia (HGPIN), low testosterone correlates with an increased risk of prostate cancer development (García-Cruz et al., 2012). This suggests that decreased activity of sex steroid signaling in the early phases of tumorigenesis could be implicated in cancer progression and/or the genesis of tumor cells populations escaping endocrine therapy. In this context, to link changes in basal extrusion, sex steroids and aggressiveness, it is noteworthy that particularly aggressive forms of ductal carcinoma *in situ* in breast (Baqai and Shousha, 2003) and intraductal cancers in prostate display both a growth initially restricted to the epithelial compartment and defects in sex steroid signaling (Zhao et al., 2018). Thus, lack of sex steroid signaling could potentially induce progression and escape to sex steroid deprivation in some endocrine-dependent forms of cancer.

Overall, we show *in vivo* that epithelial basal extrusion is a complex phenomenon: cells can extrude basally from the epithelium but still remain in the epithelial compartment. We show that repression of sex steroid signaling in an oncogenic context induces a specific change in tumor cell fate, in which these cells grow in a newly unveiled intrabasal compartment. We show that, for tumor cells of the drosophila accessory gland, this represents a key event in the genesis of tumors escaping sex steroid deprivation. Epithelial basal extrusion could therefore be a key event in tumor escape and progression, depending on activity of specific pathways such as sex steroid signaling.

## Materials and Methods

### Fly stocks and experimental crosses

y,w,HS:flp122/+;Act:FRTstopFRTGal4, UAS:GFP/CyO flies allowed conditional clonal expression of GFP. Combination with UAS:EgfrλTop (#59843) allowed conditional clonal co-expression of GFP and Egfrλtop (EGFRλ flies). EGFRλ flies were then crossed with following stocks to realize experiments: UAS-GFP.nls (#4775, control condition), UAS-EcR RNAi (#50712), UAS-Shd RNAi (#67456), UAS-Hr3 RNAi (#27253), UAS-Hr4 RNAi (#54803), UAS-Sad RNAi (#67352), UAS-Phm RNAi (#65017), UAS-Tub60D RNAi (#64856) from the Bloomington Stock Center.

### Conditional expression induction

Briefly, flippase (flp)/FRT system was activated by a 12 min heat-shock induction at 37°C during the pupal stage, to create an average of 4–6 clones per accessory gland (≈1% of total number of epithelial cells). Flies were then kept at 27 °C until the end of pupal stage. Males were collected at emergence from pupae 3 to 3.5 days after heat shock and kept for another 1 to 5 days at 27 °C before dissection.

### Immunohistochemistry and imaging

Accessory glands were dissected in PBS, fixed for 10 min in 4% paraformaldehyde, washed and permeabilized for 15 min in PBS containing 0.2% Triton (PBS-T). Glands were blocked for one hour with 0.5% of BSA in PBS-T then incubated overnight at 4°C with primary antibodies diluted in the same blocking solution. After three washes in PBS-T, glands were incubated in secondary antibody diluted 1:1000 in blocking solution for 1 hour at room temperature with DAPI (DiAminidoPhenylIndol, D8417, Sigma) 1:1000 (DNA staining) and/or Alexa633-phalloidin (A22284, Life Technology) 1:5000 (to reveal F-actin). Glands were then washed twice in PBS-T, once in PBS and subsequently mounted in Vectashield (#-1000, Vector Laboratories). Imaging was realized on Leica SP8 confocal microscope, and image stacks were processed either in ImageJ or Imaris software. All stainings were repeated a minimum of 3 times on independent experiments.

List of antibodies: Mouse Coracle (1:300, #C566.9 DSHB), mouse Fasciclin III (1:400, #7G10 DSHB), goat GFP (1:1000, #5450 Abcam), rabbit pSrc (1:500, #44-660G Invitrogen), rabbit pH3 (1:500, #06-570, Merck Millipore), rabbit cleaved-caspase 3 (1:400, #9661S, Cell Signaling), secondary antibodies coupled to different fluorophores 488 (1:1000, A11055 Invitrogen), Cy3 or Cy5 (1:1000, 711-165-152, 715-165-151, 715-175-150, Jackson Immunology).

### Clones sizes, cells and nuclei size and numbers, proliferation rate

Clones/tumors sizes, cells and nuclei volumes and numbers were determined from 3D reconstruction and automatic quantification in Imaris software. For each clone/tumor, average cell size was determined by ratio between clone size and number of nuclei in the considered clone. Proliferation rate was calculated compared to number of cells in a clone expressing GFP only (as in Fig. S3A). A typical clone is composed of 2 cells, and so 2 cells was taken as 100% proliferation. See Table 2 for the number of experiments and total numbers of individual flies that were processed for each experiment.

**Table 2.** Numbers of independent experiments (N) and individuals (n) used in the different experiments.

| Figure panel | N = Number of independent experiments | n = total number of individuals |
| --- | --- | --- |
| **Fig. 1 and 2** | N = 13 independent cohorts | n = 1140 |
| **Fig. 3M-O** | EGFRλ CTL: 6; EGFRλ EcR-KD: 8 | EGFRλ CTL: 29; EGFRλ EcR-KD: 35 |
| **Fig. 3P** | for respective column: 4, 3, 30, 4, 3, 36 | for respective column: 60, 131, 1144, 217, 123, 1525 |
| **Fig. 3Q** | EGFRλ CTL: 30; EGFRλ EcR-KD: 36 | EGFRλ CTL: 1144 EGFRλ EcR-KD: 1525 |
| **Fig. 3V** | 3 | for respective column : 32, 36, 7, 17, 22, 25 |
| **Fig. 5I** | 3 | for respective column: 203, 196, 217, 268 |
| **Fig. 5J-L** | 3 | for respective column of each panel: 53, 35, 30 |
| **Fig. 6A** | For respective column, 30, 36, 7, 9, 8, 33, 12. | For respective column: 1144, 1525, 228, 287, 207, 1618, 637. |
| **Fig. 6B** | N = 3 for all columns | For respective column: 203, 217, 52, 126, 62, 252, 167. |
| **Fig. 6J** | Left panel: 30, 14; right panel: 3 or more | Left panel: 1576, 354; right panel: 248, 130 |
| **Fig. 6K** | 3 or more | for respective column: 51, 30, 25 |

### Extra-epithelial or intrabasal tumor frequency

Extra-epithelial tumor frequency was determined as the percentage of flies that displayed at least one tumor on their accessory glands at dissection. For intrabasal tumors, individual glands were considered. See Table 2 for the number of experiments and total numbers of individual flies that were processed for each experiment.

### RNA FISH

Single-molecule RNA fluorescence in situ hybridization (smRNA-FISH) for transcripts of EcR from the accessory gland was performed using Stellaris probes (Biosearch Technologies). Probe sequences are listed in table S1. Accessory gland were dissected in PBS and fixed in 4% formaldehyde, 0.3% Triton X-100, 1x PBS for 20 min at room temperature, rinsed and permeabilized in 70% ethanol at 4°C overnight. Glands were rehydrated in smRNA FISH wash buffer (10% formamide in 2x SSC) for 10 min, resuspended in 50μL hybridization buffer (10% dextran sulphate, 10% formamide in 2xSSC) supplemented with 1.5μL of smRNA FISH probes. Hybridization was performed at 37°C overnight. Glands were then washed twice with smRNA FISH wash buffer at 37°C for 30 min and twice with 2xSSC solution. Then, DNA was stained with DAPI (1/500 dilution in 2x SSC) at room temperature for 20 min. Accessory glands were then mounted in 30μL Vectashield prior to imaging. The resulting images were processed using FIJI/ImageJ.

### Statistical analyses and human data

All experiments were repeated independently a minimum of three times (N: number of independent experiments) on numerous glands (n: number of pairs of glands or number of imaged and quantified glands for tumor size and nuclei number quantification; see Table 2). Statistical analyses were performed using GraphPad Prism 6 for tumor size and nuclei number quantification were compared by Krustal-Wallis test; % of glands with extra-glandular tumors were compared by Chi2 test; gene expression in qPCR was compared by two-tailed impaired T-test. Human data processing was done using the tools developed by Theurillat et al. and available at prostatecanceratlas.org. Expression data and pseudotime progression were individually retrieved for each considered gene using the Explore/bulk RNA function, and were grouped by stage (NORMAL, 173 samples, PRIMARY, 708 samples, mCRPC, 484 samples). Statistical analyses were then performed using GraphPad Prism 6 and quantifications were compared by Mann-Whitney test.

In all figures, significance is described as follows : * p<0.05; ** p<0.01; *** p<0.001; **** p<0.0001.

## Acknowledgments

The authors thank N. Anglaret, L. Babkina, A. De Haze, C. Gaudichet and C. Tamisier for technical help; Dr E. Chautard and Dr A. Bruhat for scientific advices; Bloomington Drosophila Stock Center (BDSC) for providing fly stocks, and Drosophila Studies Hybridoma Bank (DSHB) for providing antibodies. Dr Krzysztof Jagla for kind gift of β3 tubulin antibody. CLermont Imagerie Confocale (CLIC) facility for support with imaging. J-P Theurillat, S. Nicastri and the Prostate Cancer Atlas team for adding a new tool upon request.

---

**Figure S1:**
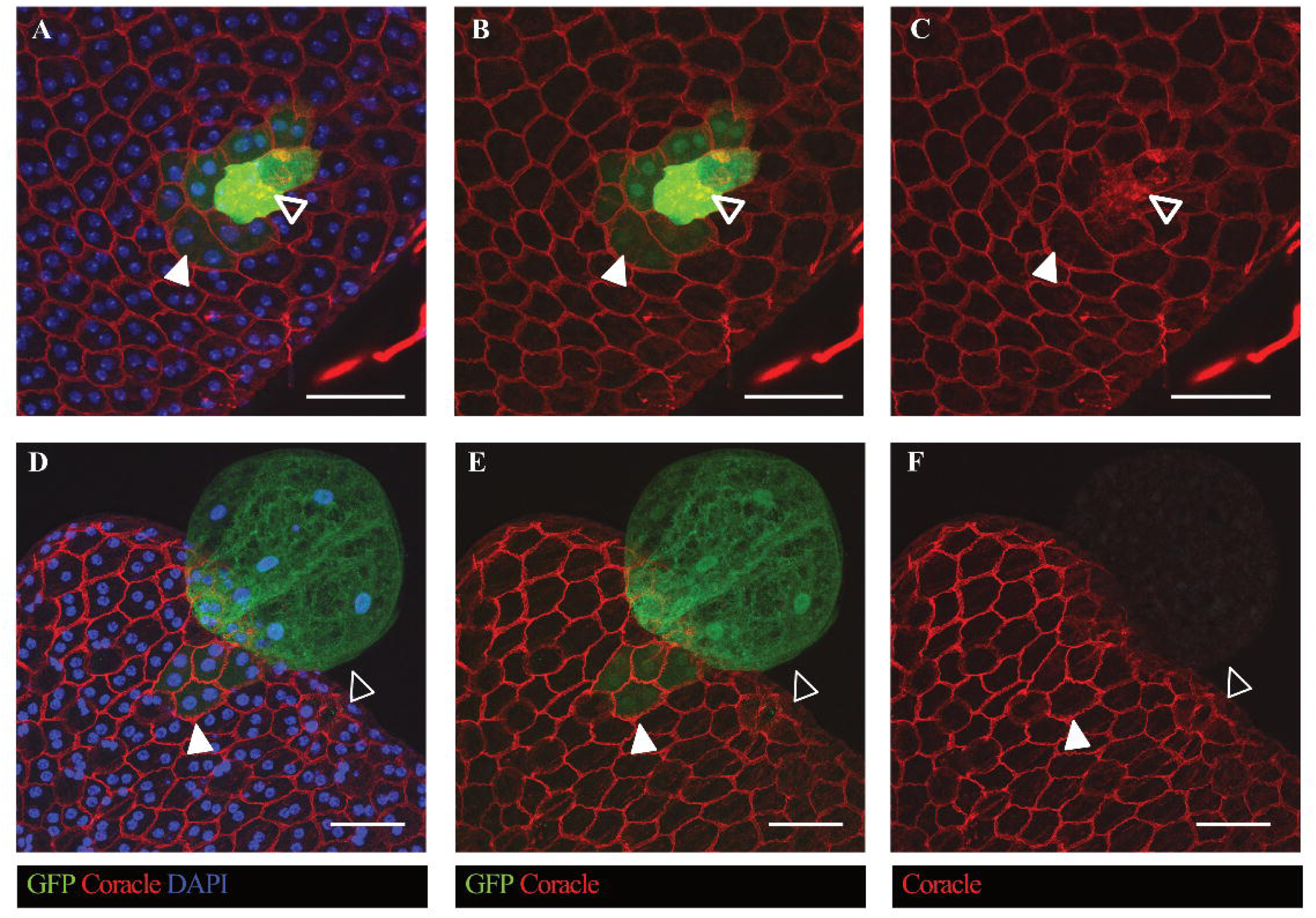
EGFRλ clonal expression induces tumorigenesis in the drosophila accessory gland. White arrowheads: mildly hyperproliferative, hypertrophic cells are well integrated into the epithelium with undisturbed cell-cell interaction (A-F, Coracle staining). Thick empty arrowheads: some of these epithelial tumor cells express higher levels of GFP (A-B), reflecting modified gene expression, and higher levels of epithelial markers. However, these markers do not properly localize at the membrane (C), indicating a loss of cohesion with neighbor cells. Empty arrowheads: some tumor cells are able to undergo epithelial basal extrusion to form extra-epithelial tumors. These cells are devoid of epithelial markers, indicating a change in their fate (F). Scale bars: 50 micrometers.

**Figure S2.**
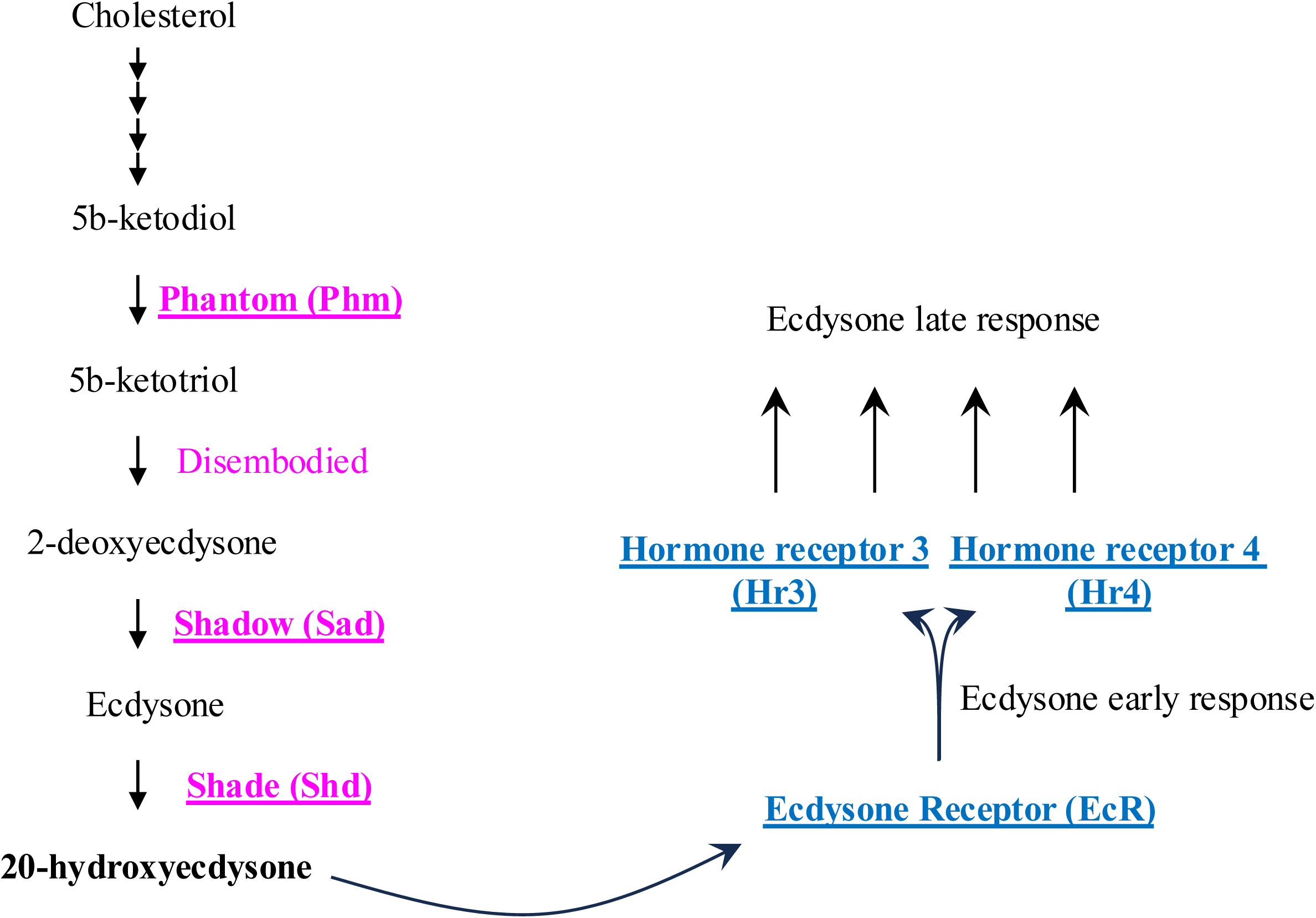
Schematic representation of the late steps of ecdysone synthesis and canonical Ecdysone Signaling. Six different genes (underlined) were independently targeted by RNAi to block tumor cell production of ecdysone (Phm, Sad), metabolization of active 20-hydroxyecdysone (Shd), or EcR Signaling (EcR, Hr3, Hr4). Results are shown in Fig. 6.

**Figure S3.**
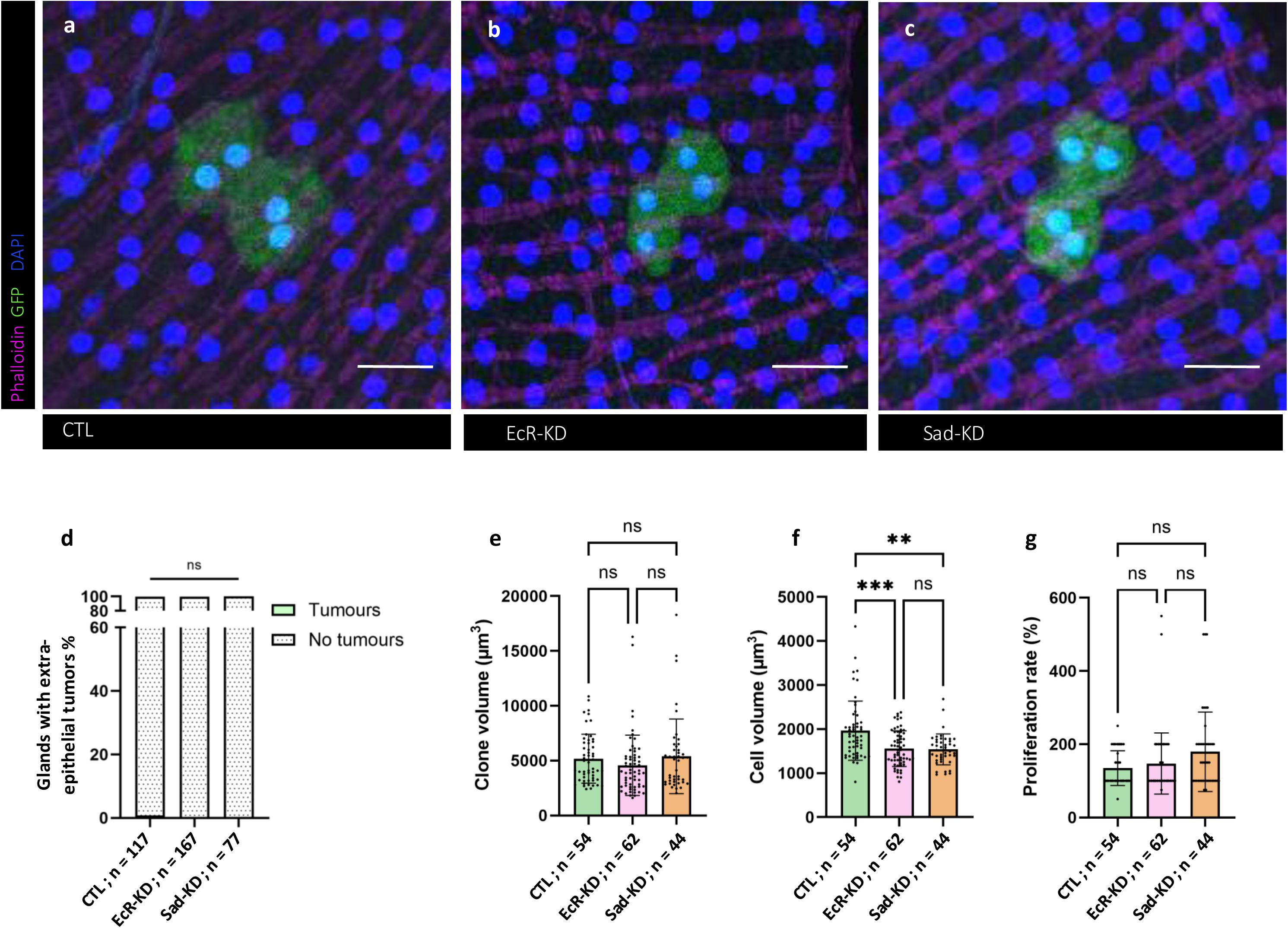
Ecdsyone deprivation does not promote tumorigenesis by itself. (a-c) In the absence of expression of an oncogene (CTL), clonal induction of NLS-GFP (a) EcR RNAi (b) or Sad RNAi (c) brings no evident phenotype to the clonal cells. (d) It does not promote tumorigenesis. (e-g) In the absence of average clone volume (e) or difference in proliferation rate compared to control (g), the only noticeable phenotype is a slightly decreased cell size in clones expressing EcR or Sad RNAi (f). Chi2 test; **P < 0.01; ***P < 0.001. Representative images in (a-c) from three or more experiments. Scale bars: 20 μm.

**Figure S4.**
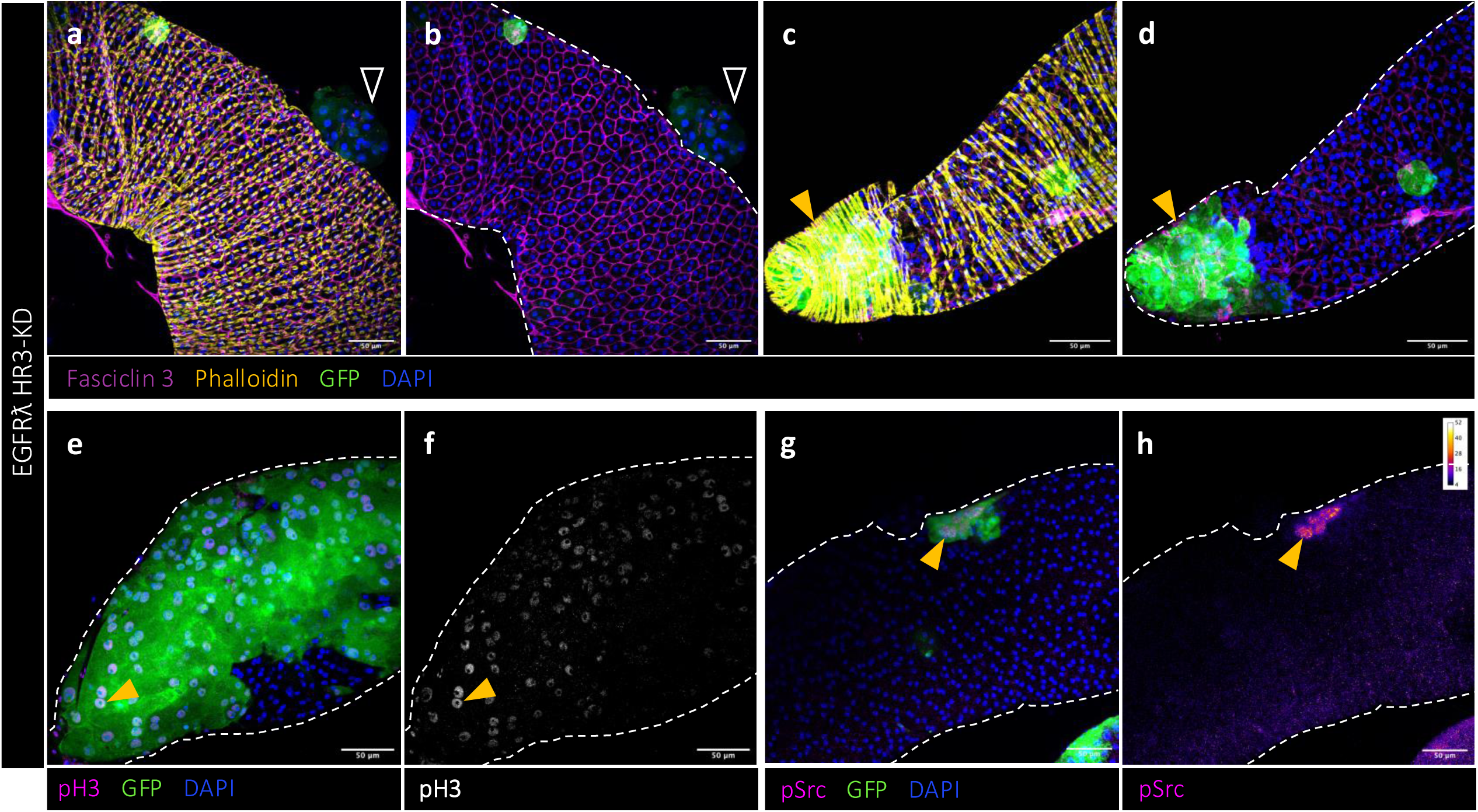
Downregulation of canonical Ecdysone Signaling induces intrabasal tumorigenesis. (a-d) Co-expression of Hr3 RNAi suppresses only partially extra-glandular tumors (empty arrowheads in (a-b)) but leads to the appearance of highly proliferative, intrabasal tumor cells into the glands (yellow arrowheads in (c-d)). (e-h) This new population displays a strong staining for classical aggressiveness markers (yellow arrowheads, pH3 staining in (e-f) and pSrc staining in (g-h)). Representative images from three or more experiments. Scale bars: 50 μM.

**Figure S5.**
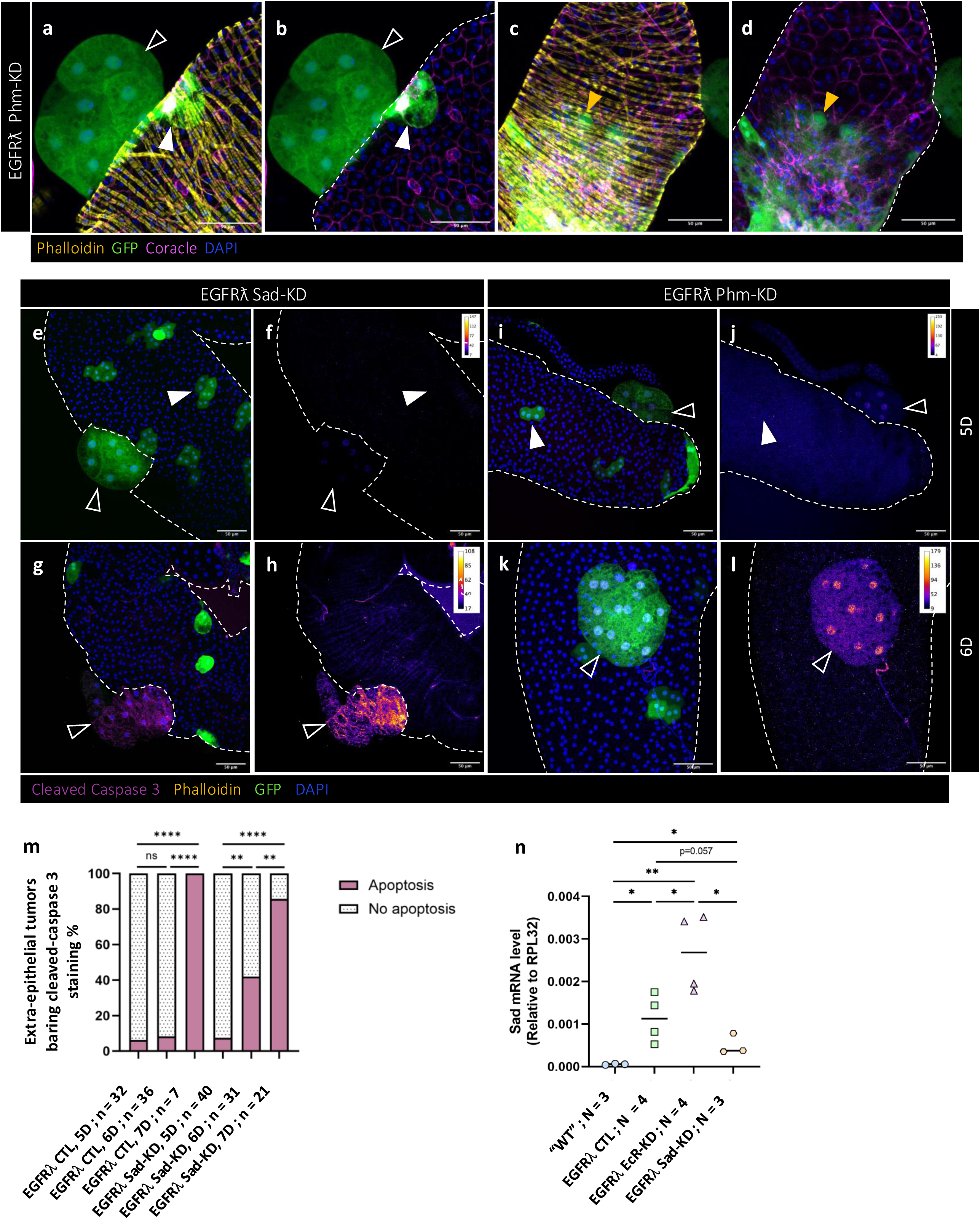
Loss of autocrine Ecdysone Signaling induces the same paradoxical response as loss of other members of the same pathway. (a-d) Co-expression of Phm RNAi suppresses only partially the formation of extra-epithelial tumors (empty arrowheads in (a-b)) but leads to the appearance of highly proliferative intrabasal tumors (yellow arrowheads in (c-d)). (e-l) Co-expression of Sad (RNAi (e-h) or Phm RNAi (i-l) induces early apoptosis of extra-epithelial tumors (at 6D, cleaved-caspase 3 staining), mimicking initial effect of steroid deprivation on human tumors (empty arrowheads). (m) Quantification of this early apoptosis phenomenon appearing at 6D for the coexpression of Sad RNAi. (n) qPCR quantification of whole glands expression of Sad for the given genotypes. “WT”: glands baring the EGFRλ genetic background, but unable to produce clones, i.e. “wild type” glands. Compared to these, oncogene induction (column 2) induces a significant overexpression of Sad mRNA; co-expression of EcR RNAi (column 3) still significantly increases Sad mRNA expression. Compared to this condition, co-expression of Sad RNAi (column 4) significantly decreases Sad mRNA expression, as expected. Representative images in (a-l) from three or more experiments. Scale bars: 50 μM.

**Figure S6.**
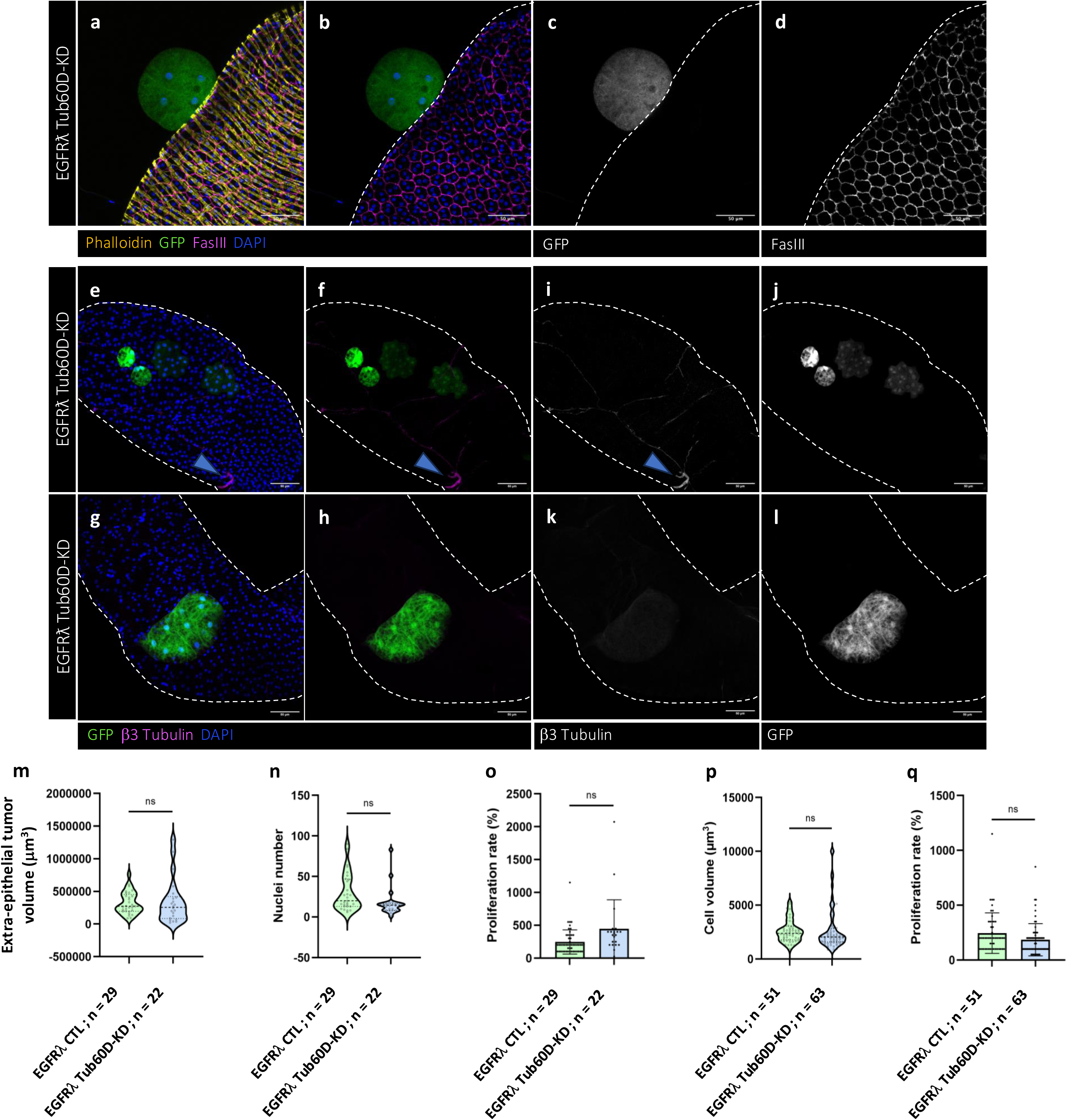
Efficient downregulation of Tub60D has little effect on epithelial tumor cells. (a-d) Co-expression of Tub60D RNAi does not affect the phenotype of extra-epithelial tumors. (e-l) Downregulation of Tub60D efficiently suppresses the accumulation of β3 Tubulin in clonal cells (e-h) and tumor cells (i-l). Remaining staining can be seen in other cell types such as tracheal cells (blue arrowheads in (e-i). (m-r) Characteristics of the different types of tumor cells co-expressing Tub60D RNAi. (m-q) Extraepithelial tumors co-expressing Tub60D RNAi display the same size (m), cell number (n) and proliferation rate (o) as EGFRλ CTL tumors. (pq) Intra-epithelial clones co-expressing Tub60D RNAi are composed of cells of same size (p) and display the same proliferation rate (q) as as EGFRλ CTL intra-epithelial clones. Chi^2^ test; ns non significant. Representative images in (a-l) from three or more experiments. Scale bars: 50 μM.

## References

Abida W, Cyrta J, Heller G, Prandi D, Armenia J, Coleman I, Cieslik M, Benelli M, Robinson D, Van Allen EM, Sboner A, Fedrizzi T, Mosquera JM, Robinson BD, De Sarkar N, Kunju LP, Tomlins S, Wu YM, Nava Rodrigues D, Loda M, Gopalan A, Reuter VE, Pritchard CC, Mateo J, Bianchini D, Miranda S, Carreira S, Rescigno P, Filipenko J, Vinson J, Montgomery RB, Beltran H, Heath EI, Scher HI, Kantoff PW, Taplin M-E, Schultz N, DeBono JS, Demichelis F, Nelson PS, Rubin MA, Chinnaiyan AM, Sawyers CL. 2019. Genomic correlates of clinical outcome in advanced prostate cancer. Proceedings of the National Academy of Sciences of the United States of America 116:11428–11436. DOI: 10.1073/pnas.1902651116, PMID: 31061129

Andersson S, Berman DM, Jenkins EP, Russell DW. 1991. Deletion of steroid 5 alpha-reductase 2 gene in male pseudohermaphroditism. Nature 354:159–161. DOI: 10.1038/354159a0, PMID: 1944596

Anton KA, Kajita M, Narumi R, Fujita Y, Tada M. 2018. Src-transformed cells hijack mitosis to extrude from the epithelium. Nature communications 9:4695. DOI: 10.1038/s41467-018-07163-4, PMID: 30410020

Arora VK, Schenkein E, Murali R, Subudhi SK, Wongvipat J, Balbas MD, Shah N, Cai L, Efstathiou E, Logothetis C, Zheng D, Sawyers CL. 2013. Glucocorticoid receptor confers resistance to antiandrogens by bypassing androgen receptor blockade. Cell 155:1309–1322. DOI: 10.1016/j.cell.2013.11.012, PMID: 24315100

Atashzar MR, Baharlou R, Karami J, Abdollahi H, Rezaei R, Pourramezan F, Zoljalali Moghaddam SH. 2020. Cancer stem cells: A review from origin to therapeutic implications. Journal of cellular physiology 235:790–803. DOI: 10.1002/jcp.29044, PMID: 31286518

Bangi E, Ang C, Smibert P, Uzilov AV, Teague AG, Antipin Y, Chen R, Hecht C, Gruszczynski N, Yon WJ, Malyshev D, Laspina D, Selkridge I, Rainey H, Moe AS, Lau CY, Taik P, Wilck E, Bhardwaj A, Sung M, Kim S, Yum K, Sebra R, Donovan M, Misiukiewicz K, Schadt EE, Posner MR, Cagan RL. 2019. A personalized platform identifies trametinib plus zoledronate for a patient with KRAS-mutant metastatic colorectal cancer. Science advances 5:eaav6528. DOI: 10.1126/sciadv.aav6528, PMID: 31131321

Bangi E, Murgia C, Teague AGS, Sansom OJ, Cagan RL. 2016. Functional exploration of colorectal cancer genomes using Drosophila. Nature communications 7:13615. DOI: 10.1038/ncomms13615, PMID: 27897178

Baqai T, Shousha S. 2003. Oestrogen receptor negativity as a marker for high-grade ductal carcinoma in situ of the breast. Histopathology 42:440–447. DOI: 10.1046/j.1365-2559.2003.01612.x, PMID: 12713620

Barbieri CE, Baca SC, Lawrence MS, Demichelis F, Blattner M, Theurillat J-P, White TA, Stojanov P, Van Allen E, Stransky N, Nickerson E, Chae S-S, Boysen G, Auclair D, Onofrio RC, Park K, Kitabayashi N, MacDonald TY, Sheikh K, Vuong T, Guiducci C, Cibulskis K, Sivachenko A, Carter SL, Saksena G, Voet D, Hussain WM, Ramos AH, Winckler W, Redman MC, Ardlie K, Tewari AK, Mosquera JM, Rupp N, Wild PJ, Moch H, Morrissey C, Nelson PS, Kantoff PW, Gabriel SB, Golub TR, Meyerson M, Lander ES, Getz G, Rubin MA, Garraway LA. 2012. Exome sequencing identifies recurrent SPOP, FOXA1 and MED12 mutations in prostate cancer. Nature genetics 44:685–689. DOI: 10.1038/ng.2279, PMID: 22610119

Baumgartner M, Radziwill G, Lorger M, Weiss A, Moelling K. 2008. c-Src-mediated epithelial cell migration and invasion regulated by PDZ binding site. Molecular and cellular biology 28:642–55. DOI: 10.1128/MCB.01024-07, PMID: 18039857

Beltran H, Prandi D, Mosquera JM, Benelli M, Puca L, Cyrta J, Marotz C, Giannopoulou E, Chakravarthi BVSK, Varambally S, Tomlins SA, Nanus DM, Tagawa ST, Van Allen EM, Elemento O, Sboner A, Garraway LA, Rubin MA, Demichelis F. 2016. Divergent clonal evolution of castration-resistant neuroendocrine prostate cancer. Nature medicine 22:298–305. DOI: 10.1038/nm.4045, PMID: 26855148

Bhatia-Gaur R, Donjacour AA, Sciavolino PJ, Kim M, Desai N, Young P, Norton CR, Gridley T, Cardiff RD, Cunha GR, Abate-Shen C, Shen MM. 1999. Roles for Nkx3.1 in prostate development and cancer. Genes & development 13:966–977. DOI: 10.1101/gad.13.8.966, PMID: 10215624

Bogden AE, Haskell PM, LePage DJ, Kelton DE, Cobb WR, Esber HJ. 1979. Growth of human tumor xenografts implanted under the renal capsule of normal immunocompetent mice. Experimental cell biology 47:281–293. DOI: 10.1159/000162947, PMID: 467773

Bolis M, Bossi D, Vallerga A, Ceserani V, Cavalli M, Impellizzieri D, Di Rito L, Zoni E, Mosole S, Elia AR, Rinaldi A, Pereira Mestre R, D’Antonio E, Ferrari M, Stoffel F, Jermini F, Gillessen S, Bubendorf L, Schraml P, Calcinotto A, Corey E, Moch H, Spahn M, Thalmann G, Kruithof-de Julio M, Rubin MA, Theurillat J-PP. 2021. Dynamic prostate cancer transcriptome analysis delineates the trajectory to disease progression. Nature communications 12:7033. DOI: 10.1038/s41467-021-26840-5, PMID: 34857732

Boysen G, Barbieri CE, Prandi D, Blattner M, Chae S-S, Dahija A, Nataraj S, Huang D, Marotz C, Xu L, Huang J, Lecca P, Chhangawala S, Liu D, Zhou P, Sboner A, de Bono JS, Demichelis F, Houvras Y, Rubin MA. 2015. SPOP mutation leads to genomic instability in prostate cancer. eLife 4. DOI: 10.7554/eLife.09207, PMID: 26374986

Bray F, Ferlay J, Soerjomataram I, Siegel RL, Torre LA, Jemal A. 2018. Global cancer statistics 2018: GLOBOCAN estimates of incidence and mortality worldwide for 36 cancers in 185 countries. CA: a cancer journal for clinicians 68:394–424. DOI: 10.3322/caac.21492, PMID: 30207593

Bruhat A, Tourmente S, Chapel S, Sobrier ML, Couderc JL, Dastugue B. 1990. Regulatory elements in the first intron contribute to transcriptional regulation of the beta 3 tubulin gene by 20-hydroxyecdysone in Drosophila Kc cells. Nucleic acids research 18:2861–2867. DOI: 10.1093/nar/18.10.2861, PMID: 2349088

Cai C, Chen Sen, Ng P, Bubley GJ, Nelson PS, Mostaghel EA, Marck B, Matsumoto AM, Simon NI, Wang H, Chen Shaoyong, Balk SP. 2011. Intratumoral de novo steroid synthesis activates androgen receptor in castration-resistant prostate cancer and is upregulated by treatment with CYP17A1 inhibitors. Cancer research 71:6503–6513. DOI: 10.1158/0008-5472.CAN-11-0532, PMID: 21868758

Cancer Genome Atlas Research Network TCGAR, Cancer Genome Atlas Research Network A, Ahn J, Akbani R, Ally A, Amin S, Andry CD, Annala M, Aprikian A, Armenia J, Arora A, Auman JT, Balasundaram M, Balu S, Barbieri CE, Bauer T, Benz CC, Bergeron A, Beroukhim R, Berrios M, Bivol A, Bodenheimer T, Boice L, Bootwalla MS, Borges dos Reis R, Boutros PC, Bowen J, Bowlby R, Boyd J, Bradley RK, Breggia A, Brimo F, Bristow CA, Brooks D, Broom BM, Bryce AH, Bubley G, Burks E, Butterfield YSN, Button M, Canes D, Carlotti CG, Carlsen R, Carmel M, Carroll PR, Carter SL, Cartun R, Carver BS, Chan JM, Chang MT, Chen Y, Cherniack AD, Chevalier S, Chin L, Cho J, Chu A, Chuah E, Chudamani S, Cibulskis K, Ciriello G, Clarke A, Cooperberg MR, Corcoran NM, Costello AJ, Cowan J, Crain D, Curley E, David K, Demchok JA, Demichelis F, Dhalla N, Dhir R, Doueik A, Drake B, Dvinge H, Dyakova N, Felau I, Ferguson ML, Frazer S, Freedland S, Fu Y, Gabriel SB, Gao J, Gardner J, Gastier-Foster JM, Gehlenborg N, Gerken M, Gerstein MB, Getz G, Godwin AK, Gopalan A, Graefen M, Graim K, Gribbin T, Guin R, Gupta M, Hadjipanayis A, Haider S, Hamel L, Hayes DN, Heiman DI, Hess J, Hoadley KA, Holbrook AH, Holt RA, Holway A, Hovens CM, Hoyle AP, Huang M, Hutter CM, Ittmann M, Iype L, Jefferys SR, Jones CD, Jones SJM, Juhl H, Kahles A, Kane CJ, Kasaian K, Kerger M, Khurana E, Kim J, Klein RJ, Kucherlapati R, Lacombe L, Ladanyi M, Lai PH, Laird PW, Lander ES, Latour M, Lawrence MS, Lau K, LeBien T, Lee D, Lee S, Lehmann K-V, Leraas KM, Leshchiner I, Leung R, Libertino JA, Lichtenberg TM, Lin P, Linehan WM, Ling S, Lippman SM, Liu J, Liu W, Lochovsky L, Loda M, Logothetis C, Lolla L, Longacre T, Lu Y, Luo J, Ma Y, Mahadeshwar HS, Mallery D, Mariamidze A, Marra MA, Mayo M, McCall S, McKercher G, Meng S, Mes-Masson A-M, Merino MJ, Meyerson M, Mieczkowski PA, Mills GB, Shaw KRM, Minner S, Moinzadeh A, Moore RA, Morris S, Morrison C, Mose LE, Mungall AJ, Murray BA, Myers JB, Naresh R, Nelson J, Nelson MA, Nelson PS, Newton Y, Noble MS, Noushmehr H, Nykter M, Pantazi A, Parfenov M, Park PJ, Parker JS, Paulauskis J, Penny R, Perou CM, Piché A, Pihl T, Pinto PA, Prandi D, Protopopov A, Ramirez NC, Rao A, Rathmell WK, Rätsch G, Ren X, Reuter VE, Reynolds SM, Rhie SK, Rieger-Christ K, Roach J, Robertson AG, Robinson B, Rubin MA, Saad F, Sadeghi S, Saksena G, Saller C, Salner A, Sanchez-Vega F, Sander C, Sandusky G, Sauter G, Sboner A, Scardino PT, Scarlata E, Schein JE, Schlomm T, Schmidt LS, Schultz N, Schumacher SE, Seidman J, Neder L, Seth S, Sharp A, Shelton C, Shelton T, Shen H, Shen R, Sherman M, Sheth M, Shi Y, Shih J, Shmulevich I, Simko J, Simon R, Simons JV, Sipahimalani P, Skelly T, Sofia HJ, Soloway MG, Song X, Sorcini A, Sougnez C, Stepa S, Stewart C, Stewart J, Stuart JM, Sullivan TB, Sun C, Sun H, Tam A, Tan D, Tang J, Tarnuzzer R, Tarvin K, Taylor BS, Teebagy P, Tenggara I, Têtu B, Tewari A, Thiessen N, Thompson T, Thorne LB, Tirapelli DP, Tomlins SA, Trevisan FA, Troncoso P, True LD, Tsourlakis MC, Tyekucheva S, Van Allen E, Van Den Berg DJ, Veluvolu U, Verhaak R, Vocke CD, Voet D, Wan Y, Wang Q, Wang W, Wang Z, Weinhold N, Weinstein JN, Weisenberger DJ, Wilkerson MD, Wise L, Witte J, Wu C-C, Wu J, Wu Y, Xu AW, Yadav SS, Yang LL, Yang LL, Yau C, Ye H, Yena P, Zeng T, Zenklusen JC, Zhang H, Zhang JJ, Zhang JJ, Zhang W, Zhong Y, Zhu K, Zmuda E, Abeshouse A, Ahn J, Akbani R, Ally A, Amin S, Andry CD, Annala M, Aprikian A, Armenia J, Arora A, Auman JT, Balasundaram M, Balu S, Barbieri CE, Bauer T, Benz CC, Bergeron A, Beroukhim R, Berrios M, Bivol A, Bodenheimer T, Boice L, Bootwalla MS, Borges dos Reis R, Boutros PC, Bowen J, Bowlby R, Boyd J, Bradley RK, Breggia A, Brimo F, Bristow CA, Brooks D, Broom BM, Bryce AH, Bubley G, Burks E, Butterfield YSN, Button M, Canes D, Carlotti CG, Carlsen R, Carmel M, Carroll PR, Carter SL, Cartun R, Carver BS, Chan JM, Chang MT, Chen Y, Cherniack AD, Chevalier S, Chin L, Cho J, Chu A, Chuah E, Chudamani S, Cibulskis K, Ciriello G, Clarke A, Cooperberg MR, Corcoran NM, Costello AJ, Cowan J, Crain D, Curley E, David K, Demchok JA, Demichelis F, Dhalla N, Dhir R, Doueik A, Drake B, Dvinge H, Dyakova N, Felau I, Ferguson ML, Frazer S, Freedland S, Fu Y, Gabriel SB, Gao J, Gardner J, Gastier-Foster JM, Gehlenborg N, Gerken M, Gerstein MB, Getz G, Godwin AK, Gopalan A, Graefen M, Graim K, Gribbin T, Guin R, Gupta M, Hadjipanayis A, Haider S, Hamel L, Hayes DN, Heiman DI, Hess J, Hoadley KA, Holbrook AH, Holt RA, Holway A, Hovens CM, Hoyle AP, Huang M, Hutter CM, Ittmann M, Iype L, Jefferys SR, Jones CD, Jones SJM, Juhl H, Kahles A, Kane CJ, Kasaian K, Kerger M, Khurana E, Kim J, Klein RJ, Kucherlapati R, Lacombe L, Ladanyi M, Lai PH, Laird PW, Lander ES, Latour M, Lawrence MS, Lau K, LeBien T, Lee D, Lee S, Lehmann K-V, Leraas KM, Leshchiner I, Leung R, Libertino JA, Lichtenberg TM, Lin P, Linehan WM, Ling S, Lippman SM, Liu J, Liu W, Lochovsky L, Loda M, Logothetis C, Lolla L, Longacre T, Lu Y, Luo J, Ma Y, Mahadeshwar HS, Mallery D, Mariamidze A, Marra MA, Mayo M, McCall S, McKercher G, Meng S, Mes-Masson A-M, Merino MJ, Meyerson M, Mieczkowski PA, Mills GB, Shaw KRM, Minner S, Moinzadeh A, Moore RA, Morris S, Morrison C, Mose LE, Mungall AJ, Murray BA, Myers JB, Naresh R, Nelson J, Nelson MA, Nelson PS, Newton Y, Noble MS, Noushmehr H, Nykter M, Pantazi A, Parfenov M, Park PJ, Parker JS, Paulauskis J, Penny R, Perou CM, Piché A, Pihl T, Pinto PA, Prandi D, Protopopov A, Ramirez NC, Rao A, Rathmell WK, Rätsch G, Ren X, Reuter VE, Reynolds SM, Rhie SK, Rieger-Christ K, Roach J, Robertson AG, Robinson B, Rubin MA, Saad F, Sadeghi S, Saksena G, Saller C, Salner A, Sanchez-Vega F, Sander C, Sandusky G, Sauter G, Sboner A, Scardino PT, Scarlata E, Schein JE, Schlomm T, Schmidt LS, Schultz N, Schumacher SE, Seidman J, Neder L, Seth S, Sharp A, Shelton C, Shelton T, Shen H, Shen R, Sherman M, Sheth M, Shi Y, Shih J, Shmulevich I, Simko J, Simon R, Simons JV, Sipahimalani P, Skelly T, Sofia HJ, Soloway MG, Song X, Sorcini A, Sougnez C, Stepa S, Stewart C, Stewart J, Stuart JM, Sullivan TB, Sun C, Sun H, Tam A, Tan D, Tang J, Tarnuzzer R, Tarvin K, Taylor BS, Teebagy P, Tenggara I, Têtu B, Tewari A, Thiessen N, Thompson T, Thorne LB, Tirapelli DP, Tomlins SA, Trevisan FA, Troncoso P, True LD, Tsourlakis MC, Tyekucheva S, Van Allen E, Van Den Berg DJ, Veluvolu U, Verhaak R, Vocke CD, Voet D, Wan Y, Wang Q, Wang W, Wang Z, Weinhold N, Weinstein JN, Weisenberger DJ, Wilkerson MD, Wise L, Witte J, Wu C-C, Wu J, Wu Y, Xu AW, Yadav SS, Yang LL, Yang LL, Yau C, Ye H, Yena P, Zeng T, Zenklusen JC, Zhang H, Zhang JJ, Zhang JJ, Zhang W, Zhong Y, Zhu K, Zmuda E. 2015. No Title. Cell 163. DOI: 10.1016/j.cell.2015.10.025, PMID: 26544944

Cao B, Qi Y, Zhang G, Xu D, Zhan Y, Alvarez X, Guo Z, Fu X, Plymate SR, Sartor O, Zhang H, Dong Y. 2014. Androgen receptor splice variants activating the full-length receptor in mediating resistance to androgen-directed therapy. Oncotarget 5:1646–1656. DOI: 10.18632/oncotarget.1802, PMID: 24722067

Carney GE, Wade AA, Sapra R, Goldstein ES, Bender M. 1997. DHR3, an ecdysone-inducible early-late gene encoding a Drosophila nuclear receptor, is required for embryogenesis. Proceedings of the National Academy of Sciences of the United States of America 94:12024–12029. DOI: 10.1073/pnas.94.22.12024, PMID: 9342356

Chountala M, Vakaloglou KM, Zervas CG. 2012. Parvin overexpression uncovers tissue-specific genetic pathways and disrupts F-actin to induce apoptosis in the developing epithelia in Drosophila. PloS one 7:e47355. DOI: 10.1371/journal.pone.0047355, PMID: 23077599

Clarke R, Brünner N, Katzenellenbogen BS, Thompson EW, Norman MJ, Koppi C, Paik S, Lippman ME, Dickson RB. 1989. Progression of human breast cancer cells from hormone-dependent to hormone-independent growth both in vitro and in vivo. Proceedings of the National Academy of Sciences of the United States of America 86:3649–3653. DOI: 10.1073/pnas.86.10.3649, PMID: 2726742

Dowsett M, Smith IE, Ebbs SR, Dixon JM, Skene A, Griffith C, Boeddinghaus I, Salter J, Detre S, Hills M, Ashley S, Francis S, Walsh G, A’Hern R. 2006. Proliferation and apoptosis as markers of benefit in neoadjuvant endocrine therapy of breast cancer. Clinical cancer research : an official journal of the American Association for Cancer Research 12:1024s–1030s. DOI: 10.1158/1078-0432.CCR-05-2127, PMID: 16467120

Dupree JM, Langille GM, Khera M, Lipshultz LI. 2014. The safety of testosterone supplementation therapy in prostate cancer. Nature reviews. Urology 11:526–530. DOI: 10.1038/nrurol.2014.163, PMID: 25069737

Ganier O, Schnerch D, Nigg EA. 2018. Structural centrosome aberrations sensitize polarized epithelia to basal cell extrusion. Open Biology 8:180044. DOI: 10.1098/rsob.180044, PMID: 29899122

García-Cruz E, Piqueras M, Ribal MJ, Huguet J, Serapiao R, Peri L, Izquierdo L, Alcaraz A. 2012. Low testosterone level predicts prostate cancer in re-biopsy in patients with high grade prostatic intraepithelial neoplasia. BJU international 110:E199–202. DOI: 10.1111/j.1464-410X.2011.10876.x, PMID: 22257176

Gauhar Z, Sun LV, Hua S, Mason CE, Fuchs F, Li T-R, Boutros M, White KP. 2009. Genomic mapping of binding regions for the Ecdysone receptor protein complex. Genome research 19:1006–1013. DOI: 10.1101/gr.081349.108, PMID: 19237466

Geller J, Albert J, Loza D, Geller S, Stoeltzing W, de la Vega D. 1978. DHT concentrations in human prostate cancer tissue. The Journal of Clinical Endocrinology and Metabolism 46:440–444. DOI: 10.1210/jcem-46-3-440, PMID: 87401

Hanahan D, Weinberg RA. 2011. Hallmarks of cancer: the next generation. Cell 144:646–674. DOI: 10.1016/j.cell.2011.02.013, PMID: 21376230

Hay CW, McEwan IJ. 2012. The impact of point mutations in the human androgen receptor: classification of mutations on the basis of transcriptional activity. PloS one 7:e32514. DOI: 10.1371/journal.pone.0032514, PMID: 22403669

Humphrey PA. 2012. Histological variants of prostatic carcinoma and their significance. Histopathology 60:59–74. DOI: 10.1111/j.1365-2559.2011.04039.x

Isaacs JT, Furuya Y, Berges R. 1994. The role of androgen in the regulation of programmed cell death/apoptosis in normal and malignant prostatic tissue. Seminars in cancer biology 5:391–400. PMID: 7849267

Ito S, Ueda T, Ueno A, Nakagawa H, Taniguchi H, Kayukawa N, Miki T. 2014. A genetic screen in Drosophila for regulators of human prostate cancer progression. Biochemical and Biophysical Research Communications 451:548–555. DOI: 10.1016/j.bbrc.2014.08.015, PMID: 25117438

Iwasa Y, Mizokami A, Miwa S, Koshida K, Namiki M. 2007. Establishment and characterization of androgen-independent human prostate cancer cell lines, LN-REC4 and LNCaP-SF, from LNCaP. International journal of urology : official journal of the Japanese Urological Association 14:233–239. DOI: 10.1111/j.1442-2042.2007.01532.x, PMID: 17430262

Kajiwara K, Chen P-K, Abe Y, Okuda S, Kon S, Adachi J, Tomonaga T, Fujita Y, Okada M. 2022. Src activation in lipid rafts confers epithelial cells with invasive potential to escape from apical extrusion during cell competition. Current biology : CB 32:3460–3476.e6. DOI: 10.1016/j.cub.2022.06.038, PMID: 35809567

Kumar A, Coleman I, Morrissey C, Zhang X, True LD, Gulati R, Etzioni R, Bolouri H, Montgomery B, White T, Lucas JM, Brown LG, Dumpit RF, DeSarkar N, Higano C, Yu EY, Coleman R, Schultz N, Fang M, Lange PH, Shendure J, Vessella RL, Nelson PS. 2016. Substantial interindividual and limited intraindividual genomic diversity among tumors from men with metastatic prostate cancer. Nature medicine 22:369–378. DOI: 10.1038/nm.4053, PMID: 26928463

Kuukasjärvi T, Kononen J, Helin H, Holli K, Isola J. 1996. Loss of estrogen receptor in recurrent breast cancer is associated with poor response to endocrine therapy. Journal of clinical oncology : official journal of the American Society of Clinical Oncology 14:2584–2589. DOI: 10.1200/JCO.1996.14.9.2584, PMID: 8823339

Labrecque MP, Coleman IM, Brown LG, True LD, Kollath L, Lakely B, Nguyen HM, Yang YC, da Costa RMG, Kaipainen A, Coleman R, Higano CS, Yu EY, Cheng HH, Mostaghel EA, Montgomery B, Schweizer MT, Hsieh AC, Lin DW, Corey E, Nelson PS, Morrissey C. 2019. Molecular profiling stratifies diverse phenotypes of treatment-refractory metastatic castration-resistant prostate cancer. The Journal of clinical investigation 129:4492–4505. DOI: 10.1172/JCI128212, PMID: 31361600

Lapuk AV, Wu C, Wyatt AW, McPherson A, McConeghy BJ, Brahmbhatt S, Mo F, Zoubeidi A, Anderson S, Bell RH, Haegert A, Shukin R, Wang Y, Fazli L, Hurtado-Coll A, Jones EC, Hach F, Hormozdiari F, Hajirasouliha I, Boutros PC, Bristow RG, Zhao Y, Marra MA, Fanjul A, Maher CA, Chinnaiyan AM, Rubin MA, Beltran H, Sahinalp SC, Gleave ME, Volik SV, Collins CC. 2012. From sequence to molecular pathology, and a mechanism driving the neuroendocrine phenotype in prostate cancer. The Journal of pathology 227:286–297. DOI: 10.1002/path.4047, PMID: 22553170

Leiblich A, Marsden L, Gandy C, Corrigan L, Jenkins R, Hamdy F, Wilson C. 2012. Bone morphogenetic protein- and mating-dependent secretory cell growth and migration in the Drosophila accessory gland. Proceedings of the National Academy of Sciences 109:19292–19297. DOI: 10.1073/pnas.1214517109, PMID: 23129615

Levine BD, Cagan RL. 2016. Drosophila Lung Cancer Models Identify Trametinib plus Statin as Candidate Therapeutic. Cell reports 14:1477–1487. DOI: 10.1016/j.celrep.2015.12.105, PMID: 26832408

Locke JA, Guns ES, Lubik AA, Adomat HH, Hendy SC, Wood CA, Ettinger SL, Gleave ME, Nelson CC. 2008. Androgen levels increase by intratumoral de novo steroidogenesis during progression of castration-resistant prostate cancer. Cancer research 68:6407–6415. DOI: 10.1158/0008-5472.CAN-07-5997, PMID: 18676866

Lu-Yao GL, Albertsen PC, Moore DF, Shih W, Lin Y, DiPaola RS, Yao S-L. 2008. Survival following primary androgen deprivation therapy among men with localized prostate cancer. JAMA 300:173–181. DOI: 10.1001/jama.300.2.173, PMID: 18612114

Marshall TW, Lloyd IE, Delalande JM, Näthke I, Rosenblatt J. 2011. The tumor suppressor adenomatous polyposis coli controls the direction in which a cell extrudes from an epithelium. Molecular Biology of the Cell 22:3962–3970. DOI: 10.1091/mbc.E11-05-0469, PMID: 21900494

Melo S, Guerrero P, Moreira Soares M, Bordin JR, Carneiro F, Carneiro P, Dias MB, Carvalho J, Figueiredo J, Seruca R, Travasso RDM. 2023. The ECM and tissue architecture are major determinants of early invasion mediated by E-cadherin dysfunction. Communications Biology 6:1132. DOI: 10.1038/s42003-023-05482-x, PMID: 37938268

Merglen A, Schmidlin F, Fioretta G, Verkooijen HM, Rapiti E, Zanetti R, Miralbell R, Bouchardy C. 2007. Short- and long-term mortality with localized prostate cancer. Archives of internal medicine 167:1944–1950. DOI: 10.1001/archinte.167.18.1944, PMID: 17923593

Michaud JE, Billups KL, Partin AW. 2015. Testosterone and prostate cancer: an evidence-based review of pathogenesis and oncologic risk. Therapeutic advances in urology 7:378–387. DOI: 10.1177/1756287215597633, PMID: 26622322

Miyoshi Y, Uemura H, Umemoto S, Sakamaki K, Morita S, Suzuki K, Shibata Y, Masumori N, Ichikawa T, Mizokami A, Sugimura Y, Nonomura N, Sakai H, Honma S, Harada M, Kubota Y. 2014. High testosterone levels in prostate tissue obtained by needle biopsy correlate with poor-prognosis factors in prostate cancer patients. BMC cancer 14:717. DOI: 10.1186/1471-2407-14-717, PMID: 25256077

Molano-Fernández M, Hickson ID, Herranz H. 2022. Cyclin E overexpression in the Drosophila accessory gland induces tissue dysplasia. Frontiers in cell and developmental biology 10:992253. DOI: 10.3389/fcell.2022.992253, PMID: 36704199

Nagabhushan M, Miller CM, Pretlow TP, Giaconia JM, Edgehouse NL, Schwartz S, Kung HJ, de Vere White RW, Gumerlock PH, Resnick MI, Amini SB, Pretlow TG. 1996. CWR22: the first human prostate cancer xenograft with strongly androgen-dependent and relapsed strains both in vivo and in soft agar. Cancer research 56:3042–3046. PMID: 8674060

Petryk A, Warren JT, Marqués G, Jarcho MP, Gilbert LI, Kahler J, Parvy J-P, Li Y, Dauphin-Villemant C, O’Connor MB. 2003. Shade is the Drosophila P450 enzyme that mediates the hydroxylation of ecdysone to the steroid insect molting hormone 20-hydroxyecdysone. Proceedings of the National Academy of Sciences of the United States of America 100:13773–13778. DOI: 10.1073/pnas.2336088100, PMID: 14610274

Rambur A, Lours-Calet C, Beaudoin C, Buñay J, Vialat M, Mirouse V, Trousson A, Renaud Y, Lobaccaro J-MA, Baron S, Morel L, de Joussineau C. 2020. Sequential Ras/MAPK and PI3K/AKT/mTOR pathways recruitment drives basal extrusion in the prostate-like gland of Drosophila. Nature communications 11:2300. DOI: 10.1038/s41467-020-16123-w, PMID: 32385236

Rambur A, Vialat M, Beaudoin C, Lours-Calet C, Lobaccaro J-M, Baron S, Morel L, de Joussineau C. 2021. Drosophila Accessory Gland: A Complementary In Vivo Model to Bring New Insight to Prostate Cancer. Cells 10. DOI: 10.3390/cells10092387, PMID: 34572036

Read RD. 2011. Drosophila melanogaster as a model system for human brain cancers. Glia 59:1364–76. DOI: 10.1002/glia.21148, PMID: 21538561

Read RD, Cavenee WK, Furnari FB, Thomas JB. 2009. A Drosophila Model for EGFR-Ras and PI3K-Dependent Human Glioma. PLoS Genetics 5:e1000374. DOI: 10.1371/journal.pgen.1000374

Rodriguez D, Ramkairsingh M, Lin X, Kapoor A, Major P, Tang D. 2019. The Central Contributions of Breast Cancer Stem Cells in Developing Resistance to Endocrine Therapy in Estrogen Receptor (ER)-Positive Breast Cancer. Cancers 11. DOI: 10.3390/cancers11071028, PMID: 31336602

Rybak AP, Bristow RG, Kapoor A. 2015. Prostate cancer stem cells: deciphering the origins and pathways involved in prostate tumorigenesis and aggression. Oncotarget 6:1900–1919. DOI: 10.18632/oncotarget.2953, PMID: 25595909

Sekar A, Leiblich A, Wainwright SM, Mendes CC, Sarma D, Hellberg JEEU, Gandy C, Goberdhan DCI, Hamdy FC, Wilson C. 2023. Rbf/E2F1 control growth and endoreplication via steroid-independent Ecdysone Receptor signalling in Drosophila prostate-like secondary cells. PLoS genetics 19:e1010815. DOI: 10.1371/journal.pgen.1010815, PMID: 37363926

Sharma V, Pandey AK, Kumar A, Misra S, Gupta HPK, Gupta S, Singh A, Buehner NA, Ravi Ram K. 2017. Functional male accessory glands and fertility in Drosophila require novel ecdysone receptor. PLoS genetics 13:e1006788. DOI: 10.1371/journal.pgen.1006788, PMID: 28493870

Sharp A, Coleman I, Yuan W, Sprenger C, Dolling D, Rodrigues DN, Russo JW, Figueiredo I, Bertan C, Seed G, Riisnaes R, Uo T, Neeb A, Welti J, Morrissey C, Carreira S, Luo J, Nelson PS, Balk SP, True LD, de Bono JS, Plymate SR. 2019. Androgen receptor splice variant-7 expression emerges with castration resistance in prostate cancer. The Journal of clinical investigation 129:192–208. DOI: 10.1172/JCI122819, PMID: 30334814

Shen J, Lu J, Sui L, Wang D, Yin M, Hoffmann I, Legler A, Pflugfelder GO. 2014. The orthologous Tbx transcription factors Omb and TBX2 induce epithelial cell migration and extrusion in vivo without involvement of matrix metalloproteinases. Oncotarget 5:11998–12015. DOI: 10.18632/oncotarget.2426, PMID: 25344916

Shirai T, Sekai M, Kozawa K, Sato N, Tanimura N, Kon S, Matsumoto T, Murakami T, Ito S, Tilston-Lunel A, Varelas X, Fujita Y. 2022. Basal extrusion of single-oncogenic mutant cells induces dome-like structures with altered microenvironments. Cancer science 113:3710–3721. DOI: 10.1111/cas.15483, PMID: 35816400

Slattum G, McGee KM, Rosenblatt J. 2009. P115 RhoGEF and microtubules decide the direction apoptotic cells extrude from an epithelium. The Journal of Cell Biology 186:693–702. DOI: 10.1083/jcb.200903079, PMID: 19720875

Stelloo S, Nevedomskaya E, Kim Y, Schuurman K, Valle-Encinas E, Lobo J, Krijgsman O, Peeper DS, Chang SL, Feng FY-C, Wessels LFA, Henrique R, Jerónimo C, Bergman AM, Zwart W. 2018. Integrative epigenetic taxonomy of primary prostate cancer. Nature communications 9:4900. DOI: 10.1038/s41467-018-07270-2, PMID: 30464211

Suntsova M, Gaifullin N, Allina D, Reshetun A, Li X, Mendeleeva L, Surin V, Sergeeva A, Spirin P, Prassolov V, Morgan A, Garazha A, Sorokin M, Buzdin A. 2019. Atlas of RNA sequencing profiles for normal human tissues. Scientific data 6:36. DOI: 10.1038/s41597-019-0043-4, PMID: 31015567

Taniguchi K, Kokuryo A, Imano T, Nakagoshi H, Adachi-Yamada T. 2018. Binucleation of Accessory Gland Lobe Contributes to Effective Ejection of Seminal Fluid in Drosophila *melanogaster*. Zoological Science 35:446–458. DOI: 10.2108/zs170188, PMID: 30298781

The Genotype-Tissue Expression (GTEx) project. 2013. Nature genetics 45:580–585. DOI: 10.1038/ng.2653, PMID: 23715323

Turpin A, Delliaux C, Parent P, Chevalier H, Escudero-Iriarte C, Bonardi F, Vanpouille N, Flourens A, Querol J, Carnot A, Leroy X, Herranz N, Lanel T, Villers A, Olivier J, Touzet H, de Launoit Y, Tian TV, Duterque-Coquillaud M. 2023. Fascin-1 expression is associated with neuroendocrine prostate cancer and directly suppressed by androgen receptor. British Journal of Cancer 129:1903–1914. DOI: 10.1038/s41416-023-02449-x

Ventrella R, Kim SK, Sheridan J, Grata A, Bresteau E, Hassan OA, Suva EE, Walentek P, Mitchell BJ. 2023. Bidirectional multiciliated cell extrusion is controlled by Notch-driven basal extrusion and Piezo1-driven apical extrusion. Development (Cambridge, England) 150:dev201612. DOI: 10.1242/dev.201612, PMID: 37602491

Vialat M, Baabdaty E, Trousson A, Kocer A, Lobaccaro J-MA, Baron S, Morel L, de Joussineau C. 2024. Cholesterol Dietary Intake and Tumor Cell Homeostasis Drive Early Epithelial Tumorigenesis: A Potential Modelization of Early Prostate Tumorigenesis. Cancers 16. DOI: 10.3390/cancers16112153, PMID: 38893271

Villeneuve C, Lagoutte E, de Plater L, Mathieu S, Manneville J-B, Maître J-L, Chavrier P, Rossé C. 2019. aPKCi triggers basal extrusion of luminal mammary epithelial cells by tuning contractility and vinculin localization at cell junctions. Proceedings of the National Academy of Sciences of the United States of America 116:24108–24114. DOI: 10.1073/pnas.1906779116, PMID: 31699818

Wilson C, Leiblich A, Goberdhan DCI, Hamdy F. 2017. The Drosophila Accessory Gland as a Model for Prostate Cancer and Other Pathologies. Current Topics in Developmental Biology. p. 339–375. DOI: 10.1016/bs.ctdb.2016.06.001, PMID: 28057306

Wu HC, Hsieh JT, Gleave ME, Brown NM, Pathak S, Chung LW. 1994. Derivation of androgen-independent human LNCaP prostatic cancer cell sublines: role of bone stromal cells. International journal of cancer 57:406–412. DOI: 10.1002/ijc.2910570319, PMID: 8169003

Xue L, Noll M. 2002. Dual role of the Pax gene paired in accessory gland development of Drosophila. Development (Cambridge, England) 129:339–46. PMID: 11807027

Zhao J, Sun G, Zhu S, Dai J, Chen J, Zhang M, Ni Y, Zhang H, Shen P, Zhao X, Zhang B, Pan X, Nie L, Yin X, Liang J, Zhang X, Wang Z, Zhu X, Liao B, Liu Z, Armstrong CM, Gao AC, Huang H, Chen N, Zeng H, Porter LH, Hashimoto K, Lawrence MG, Pezaro C, Clouston D, Wang H, Papargiris M, Thorne H, Li J, Ryan A, Norden S, Moon D, Bolton DM, Sengupta S, Frydenberg M, Murphy DG, Risbridger GP, Taylor RA, Bonkhoff H, Wheeler TM, van der Kwast TH, Magi-Galluzzi C, Montironi R, Cohen RJ. 2018. Intraductal carcinoma of the prostate: precursor or aggressive phenotype of prostate cancer? BJU international 121:971–978. DOI: 10.1002/pros.22579, PMID: 28977728

Zhou J-R, Yu L, Zerbini LF, Libermann TA, Blackburn GL. 2004. Progression to androgen-independent LNCaP human prostate tumors: cellular and molecular alterations. International journal of cancer 110:800–806. DOI: 10.1002/ijc.20206, PMID: 15170660

Zoeller JJ, Bronson RT, Selfors LM, Mills GB, Brugge JS. 2017. Niche-localized tumor cells are protected from HER2-targeted therapy via upregulation of an anti-apoptotic program in vivo. NPJ breast cancer 3:18. DOI: 10.1038/s41523-017-0020-z, PMID: 28649658

Zundelevich A, Dadiani M, Kahana-Edwin S, Itay A, Sella T, Gadot M, Cesarkas K, Farage-Barhom S, Saar EG, Eyal E, Kol N, Pavlovski A, Balint-Lahat N, Dick-Necula D, Barshack I, Kaufman B, Gal-Yam EN. 2020. ESR1 mutations are frequent in newly diagnosed metastatic and loco-regional recurrence of endocrine-treated breast cancer and carry worse prognosis. Breast cancer research : BCR 22:16. DOI: 10.1186/s13058-020-1246-5, PMID: 32014063

